# Cell cycle dynamics controls fluidity of the developing mouse neuroepithelium

**DOI:** 10.1101/2022.01.20.477048

**Authors:** Laura Bocanegra-Moreno, Amrita Singh, Edouard Hannezo, Marcin Zagorski, Anna Kicheva

## Abstract

As organs are remodelled by morphogenetic changes and pattern formation during development, their material properties may change. To address whether and how this occurs in the mouse neural tube, we combined highly resolved mosaic analysis, biophysical modelling and perturbation experiments. We found that at early developmental stages the neuroepithelium surprisingly maintains both high junctional tension and high fluidity. This is achieved via a previously unrecognized mechanism in which interkinetic nuclear movements generate cell area dynamics that drive extensive cell rearrangements. Over time, the proliferation rate declines, effectively solidifying the tissue. Thus, unlike well-studied jamming transitions, the solidification we uncovered resembles a glass transition that depends on the dynamics of stresses generated by proliferation and differentiation. This new link between epithelial fluidity, interkinetic movements and cell cycle dynamics has implications for the precision of pattern formation and could be relevant to multiple developing tissues.

## Introduction

Cells within developing tissues reorganize at the same time as pattern formation and tissue growth take place. The extent and dynamics of cell rearrangements can change substantially throughout tissue development (*1, 2*). In most cases, these changes have been proposed to result from solid-fluid transitions driven by alterations in cell density, internal myosin and/or cadherin-mediated adhesion forces at cell junctions, or external mechanical forces (*2–7*). Cell rearrangements have also been shown in theory and in some experimental situations to depend on active stresses within tissues, such as the ones generated by cell division (*8–11*). Yet, in many cases, the dynamics of cell rearrangements and the factors that control them are poorly understood. As a consequence, the impact of cell reorganization on pattern formation and morphogenesis has remained unclear.

In the developing spinal cord, pattern formation along the dorsoventral (DV) axis occurs at the earliest stages of neural tube formation (*12*). At these stages, significant cell movements have been observed in the forming neural tube of zebrafish embryos, which undergo secondary neurulation (*13*). Short time-scale live-imaging and lineage tracing experiments (*14–20*) suggest that cell rearrangements also occur in the amniote spinal cord, which forms through primary neurulation. However, the quantitative dynamics of cell rearrangements during development, and the factors that control them, remain unclear. Here we use highly resolved clonal analysis to measure the rate of cell rearrangements in the mouse neuroepithelium over time. We determine the contribution of cell divisions and mechanical forces to epithelial dynamics and assess how rearrangements can affect pattern formation.

## Results

### Cell rearrangements decline over time

To reliably and quantitively measure cell rearrangements, we used Mosaic Analysis with Double Markers (MADM) to generate extremely sparse clonal labelling (Figure 1, (*21, 22*)). In this approach, Sox2-CreERT2-induced recombination results in the expression of red (tdTomato) and green (EGFP) fluorescent proteins, either in distinct daughter cells or in the same cell (Figures 1A, S1, see Methods). Recombination events occur with very low probability - we detected between 1 and 4 clones per spinal cord (Figure S1A).

**Figure 1.**
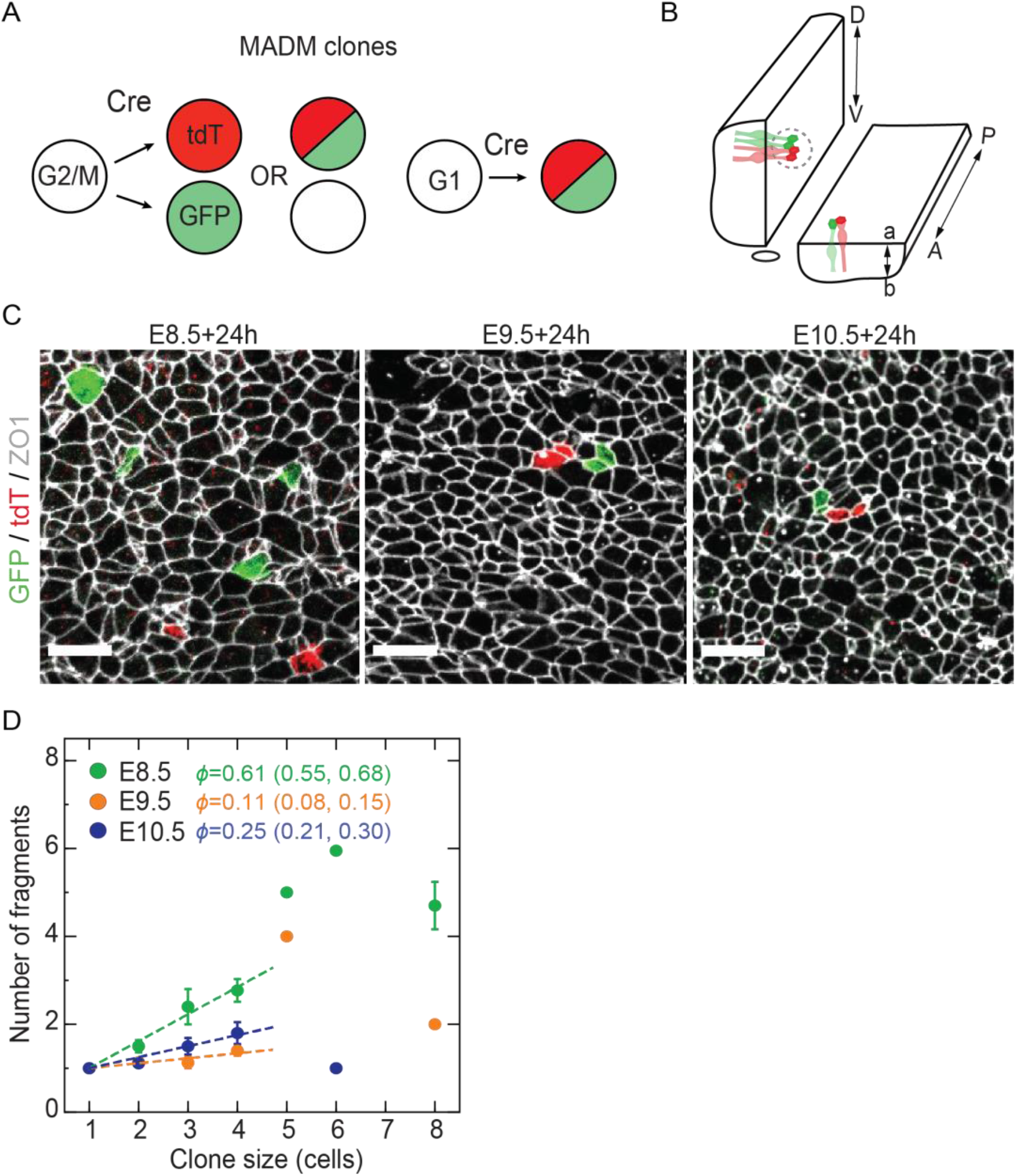
Tracing of cell rearrangements in the developing spinal cord using MADM clones. **A**. MADM labelling. **B**. Scheme of embryo mounting. **C**. MADM clones induced at the indicated stages and analysed 24h later. Scale bars, 10 μm. **D**. Mean number of fragments per clone ± SEM. Clones analysed 24h post injection. Corresponding *ϕ* with 95% CI. Sample sizes, see Table S1.

We induced MADM clonal labelling by injecting pregnant mothers with tamoxifen at embryonic days E8.5, E9.5 and E10.5, and harvested the embryos 24h later. The cytosolic localisation of the fluorescent reporters allows labelled cells to be detected at the apical surface of the epithelium (Figure 1B, C). In addition, we co-immunostained the tissue for the tight junction marker ZO1, which allowed us to segment individual cells and determine precisely the number and neighbour relationships of labelled cells.

Previous estimates showed that the net growth rate of the neuroepithelium progenitor layer declines from 0.084±0.003 h^-1^ (mean±SEM) at E8.5 to 0.041±0.002 h^-1^ at E10.5 (*23*). Consistent with these data, the mean clone sizes of MADM clones decline from 4.1±0.3 to 2.1±0.1 cells per clone, which corresponds to a growth rate of 0.087±0.009 h^-1^ and 0.046±0.004 h^-1^ at E8.5 and E10.5, respectively (Figure S2A-B, see Methods). The clone size distributions further show that 2, 4 and 8 cell clones are the most abundant in the E8.5 dataset, indicating that cell divide up to 3 times and without significant progenitor loss (Figure S2C). At E9.5 and E10.5 larger clones are progressively under-represented, consistent with the longer cell cycle length and loss of progenitors due to terminal differentiation at these stages. Together, these observations indicated that the MADM clones accurately reflect the dynamics of tissue growth.

To estimate the extent of cell rearrangements, we next analysed the clonal shapes. In many tissues, such as the Drosophila wing disc or mouse skin, uniform tissue growth with minimal cell rearrangements results in the formation of coherent clones (*24, 25*). By contrast, cell rearrangements result in clone fragmentation, whereby subsets of the labelled cells are surrounded by non-labelled neighbours. We therefore used the number of fragments per clone as a readout of cell rearrangements. To exclude effects of clone size, we measured the fragments for clones of a given size. The number of fragments per clone depends linearly on clone size for small clone sizes (≤4 cells) for which reliable statistics can be obtained (Figure 1D). This allows us to define the fragmentation coefficient *ϕ* as the slope of a linear fit to number of fragments as a function of clone size (for clone sizes ≤4 cells). We found that MADM clones labelled at E9.5 and E10.5 had very few fragments, corresponding to *ϕ* = 0.11 (0.08, 0.15 95%CI) and 0.25 (0.21, 0.30 95%CI), respectively. By contrast, clones labelled at E8.5 were highly fragmented with *ϕ* = 0.61 (0.55, 0.68 95%CI) (Figure 1D).

Consistent with their higher fragmentation, clones labelled at E8.5 had dispersed at a larger maximum distance from the clone centroid - 10.2±1.4 μm, while clones labelled at E9.5 and E10.5 dispersed up to 3.3±0.4μm and 3.0±0.8 μm, respectively (Figure S3A). The dispersal of cells was nearly isotropic with respect to clone center, with exception of clones in the pMN domain which have a larger AP/DV aspect ratio compared to clones in other domains at E10.5 of development (Figure S3B). This effect is consistent with our previous observations and is linked with the differentiation dynamics in the pMN domain (*26*). Altogether, these results indicate that cell rearrangements occur more frequently prior to E9.5 and sharply decline at later stages.

### Interkinetic nuclear movements fluidize the neuroepithelium

To investigate how the high extent of cell rearrangements at early developmental stages is achieved, we used a 2D vertex model of the apical surface of the neuroepithelium that we previously developed (*26*). A distinct feature of our model is that it takes into account not only cell division, but also the inter-kinetic nuclear movements (IKNM) during the cell cycle, which cause apical area fluctuations. Similar to other vertex models (*27*), our model has distinct behaviours as a function of normalized tension 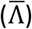 and normalized contractility 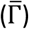 parameters. To determine model parameters that reproduce the experimentally observed clone fragmentation patterns, we performed a systematic screen in the intermediate region of 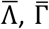 parameter space, where the network configuration is expected to be most similar to epithelial tissues (Figure 2A, see Table S2) (*27, 28*). We traced clones in silico for 16 hours, which corresponds to the duration of Cre activity in experiments (see Methods, Figures S1B, S2D). However, we observed that with this configuration, without any sources of noise, the model could not reproduce the experimentally observed clone fragmentation regardless of the rates of proliferation, cell loss, or tissue growth anisotropy (Figure S4A-C).

**Figure 2.**
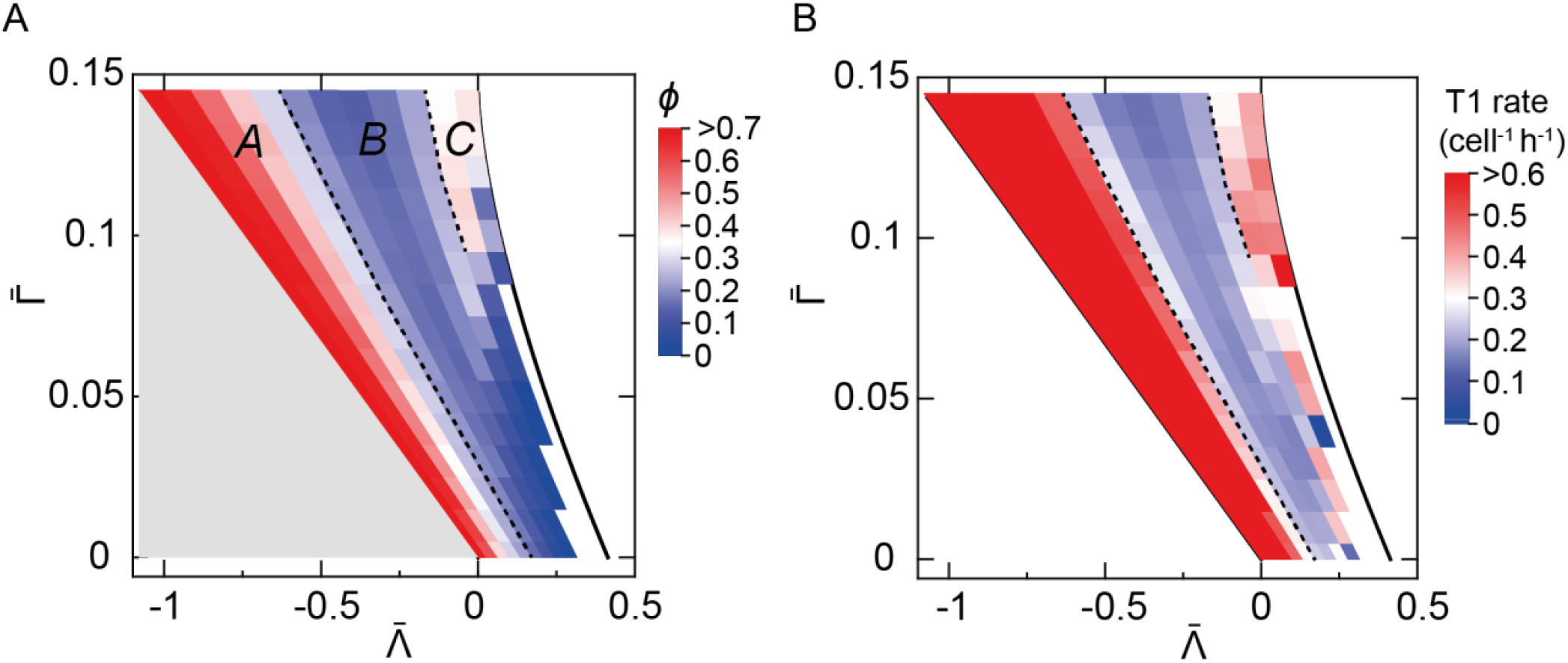
Regions with distinct clonal fragmentation in the vertex model parameter space. **A**. Coefficient of fragmentation for different values of 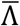 and 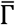, *k*_*p*_ = 0.09h^-1^, *k*_*n*_ = 0. Dashed lines correspond to *ϕ* = 0.3 and delineate regions A, B and C. Grey region, fluid ground state of the model. White region, unstable due to area collapse. **B**. Mean rate of T1 transition events (cell^-1^ h^-1^) across the 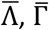 parameter space.

In other epithelia, fluctuations in the levels of myosin activity at cell junctions cause variation in the line tension and edge lengths of cells (*29*). We reasoned that a similar effect might occur in the neuroepithelium and modelled line tension fluctuations as an Ornstein-Uhlenbeck process. To this end, we introduced a noise term in 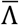, drawn from a Gaussian distribution with characteristic deviation time *σ*. This increased clonal fragmentation (Figures 2A, S4D), and in particular *σ* = 0.02 produced fragmentation coefficients similar to experimental observations.

With line tension fluctuations taken into account, the model remarkably revealed that fragmentation rates significantly vary across the 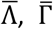 parameter space. In particular, two sharp transitions subdivide the parameter space into three subregions, which we refer to as regions A, B and C (Figure 2A). Regions A and C have high fragmentation (*ϕ* ≥ 0.3) and high T1 transition rate, while region B has low fragmentation (*ϕ* < 0.3) and T1 rate (Figure 2A-B). Previous studies of vertex models have observed a density-independent fluid-to-solid phase transition, which is characterized by a change in the cell shape index and could be analogous to the transition between regions A and B that we observe (*1, 7, 27*). By contrast, the high rate of T1 transitions in region C has not previously been observed and is surprising, given that the ground state of the model in this region is solid (*27, 28*). The high fragmentation coefficient that we observed at E8.5 (Figure 1D) is consistent with both high fragmentation regions A and C. Hence, more than one mechanism, captured by either region A or C, could explain how the high fragmentation rates are achieved at early developmental stages.

To distinguish potential mechanisms and understand how fragmentation is achieved in the E8.5 neural tube, we compared the cell shapes in simulations of regions A vs C (Figures 3, S5). Several first order descriptors of cell shape (see Table S3) were similar between regions A and C. For instance, these regions were characterized by high cell shape index and low packing order, measured by the fraction of hexagons, which are indicators of tissue fluidity (*7*) (Figure 3A-B). By contrast, a subset of cell shape descriptors differed between regions A and C. These included the coefficients of variation (CV) of the cell area, perimeter, and elongation, as well as the area ratio slope (Figures 3C, S5). The most striking difference between regions A and C was that only region C had high cell area CV, while in region A cell areas were nearly uniform (Figure 3C).

**Figure 3.**
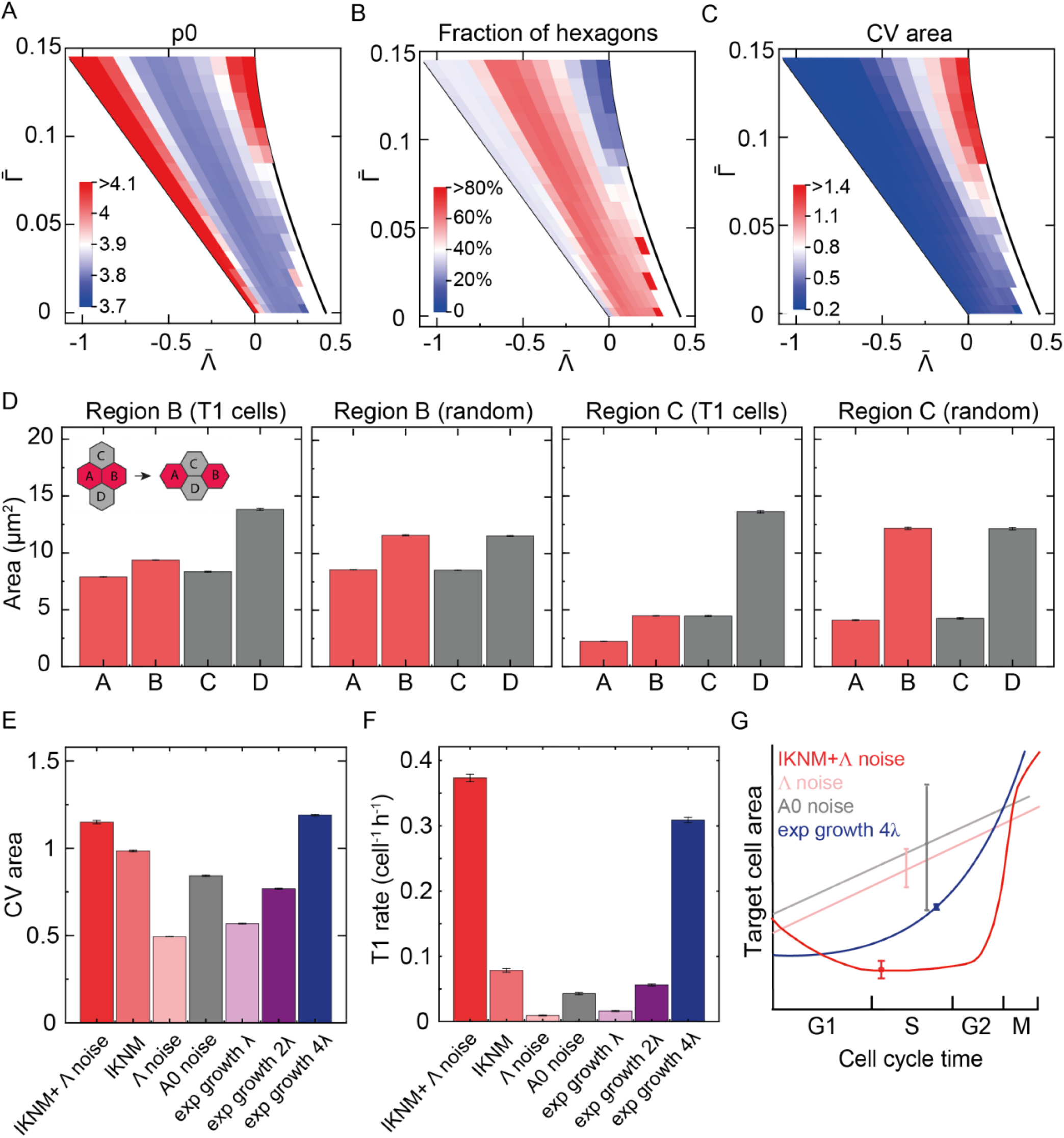
Cell area kinetics underlies extensive cell rearrangements. **A-C**. Mean cell shape index, fraction of hexagons, and coefficient of variation of apical cell areas for 10 simulations per 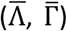 parameter set. **D**. Neighboring cells A, B, C, D undergoing T1 transition (T1 cells) or randomly assigned (random). **E, F**. Mean cell area CV (E) and T1 rate (F) for simulations with variable cell area growth during the cell cycle, n=10 simulations per condition, region 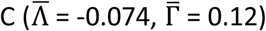. IKNM+Λ noise (default condition), IKNM (no noise in line tension), Λ noise (cell divisions without IKNM, with noise in 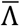), A0 noise (target area randomly drawn from a distribution), exp growth (exponential growth of the target area). Details in Table S4. **G**. Schematic of the change in mean cell area during the cell cycle in the conditions in E-F. Error bars SEM.

To test if the high cell size variability is important for clonal fragmentation in region C, we first quantified the cell areas of quartets of adjoining cells undergoing a T1 transition in silico and compared them to randomly selected quartets. This analysis showed that T1 transitions occur preferentially between cell pairs with large area discrepancy (Figure 3D). We next examined the origin of cell size heterogeneity in region C. The variation in cell areas in region C was critically dependent on the implementation of IKNM in the model and it was further amplified by the noise in line tension (Figure 3E). As expected, the implementation of IKNM and noise increased not only the cell area CV, but also the extent of T1 transitions and clone fragmentation (Figure 3F). However, noise decreased the cell shape index in region C (Figure S6), indicating that the relationship between cell rearrangement rates and cell shapes is complex when several factors, such as proliferation, IKNM and line tension noise, all generate active stresses within the tissue. Further analysis showed that in the absence of cell divisions, the rate of T1 transitions and clone fragmentation in region C was close to zero (<0.01 cell^-1^ h^-1^), as expected from the solid ground state of the network in this parameter region. This indicates that cell divisions with IKNM induce a transition that effectively fluidizes the tissue.

We reasoned that IKNM could lead to increased cell rearrangements in region C either by directly increasing the cell area heterogeneity or by dynamically altering the target area of cells during the cell cycle in a specific manner. To distinguish these possibilities, we simulated a tissue without IKNM but with target cell areas drawn from a random distribution with CV comparable to the experimentally measured one (Methods). These simulations show that increasing the target cell area heterogeneity is not sufficient to increase the rate of T1 transitions (Figure 3E-G). By contrast, apical target area that increases exponentially over the cell cycle can generate increased T1 transitions. Furthermore, the sharper the increase in exponential growth rate, the higher the area heterogeneity, and the higher the rate of T1 transitions (Figure 3E-G). Altogether, this analysis suggests that kinetics of cell area growth produced by IKNM generates dynamical changes in relative cellular area which are key for rearrangements in region C.

Having established the different mechanisms by which cell rearrangements are achieved in regions A and C, we next asked whether the rearrangements that we observe in the E8.5 neuroepithelium correspond to region A or region C. Our analysis of cell shapes in silico indicated that comparing the area CV of E8.5 neuroepithelia to the model (Figure 3C) would allow us to distinguish between these possibilities. We therefore immunostained the tight junctions in E8.5 embryos and segmented the cell shapes. This analysis revealed that the CV of cell areas at E8.5 is 0.86±0.02, which is quantitatively consistent with the values observed in region C, but not in region A (Figure 3C). This suggests that the high fluidity of the E8.5 epithelium is achieved in the regime of high junctional tension and contractility characteristic of region C.

### Cell cycle lengthening and terminal differentiation decrease neuroepithelial cell rearrangements

The impact of IKNM on clone fragmentation revealed by our model suggests that the cell division rate could be critical for controlling the extent of cell rearrangements in the neuroepithelium. Between E8.5 and E10.5 of development the proliferation rate decreases and terminal differentiation commences (*23*), which lowers the net growth rate of the tissue approximately 2-fold between (Figure S2B). This change in cell division and differentiation could represent a change in the active stresses that generate fluctuations in the tissue and in this way cause tissue solidification over time. To test this possibility, we lowered the proliferation rate in the vertex model simulations from 0.09 to 0.03 h^-1^. This resulted in a significant decline of the fragmentation coefficient of clones throughout most of the 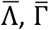 parameter space (Figures 4A, S7A). In region C, *ϕ* declined ∼2-fold and a corresponding decline in T1 rates was observed (Figure S7B), while the cell area CV was reduced to a lesser extent and remained significantly higher than in region B (Figure S7C). This reduction in the fragmentation coefficient in the model is reminiscent of the experimentally observed reduction in *ϕ* (Figure 1D), suggesting that the decreasing rate of proliferation over time is a key driver of the decline in cell rearrangements.

**Figure 4.**
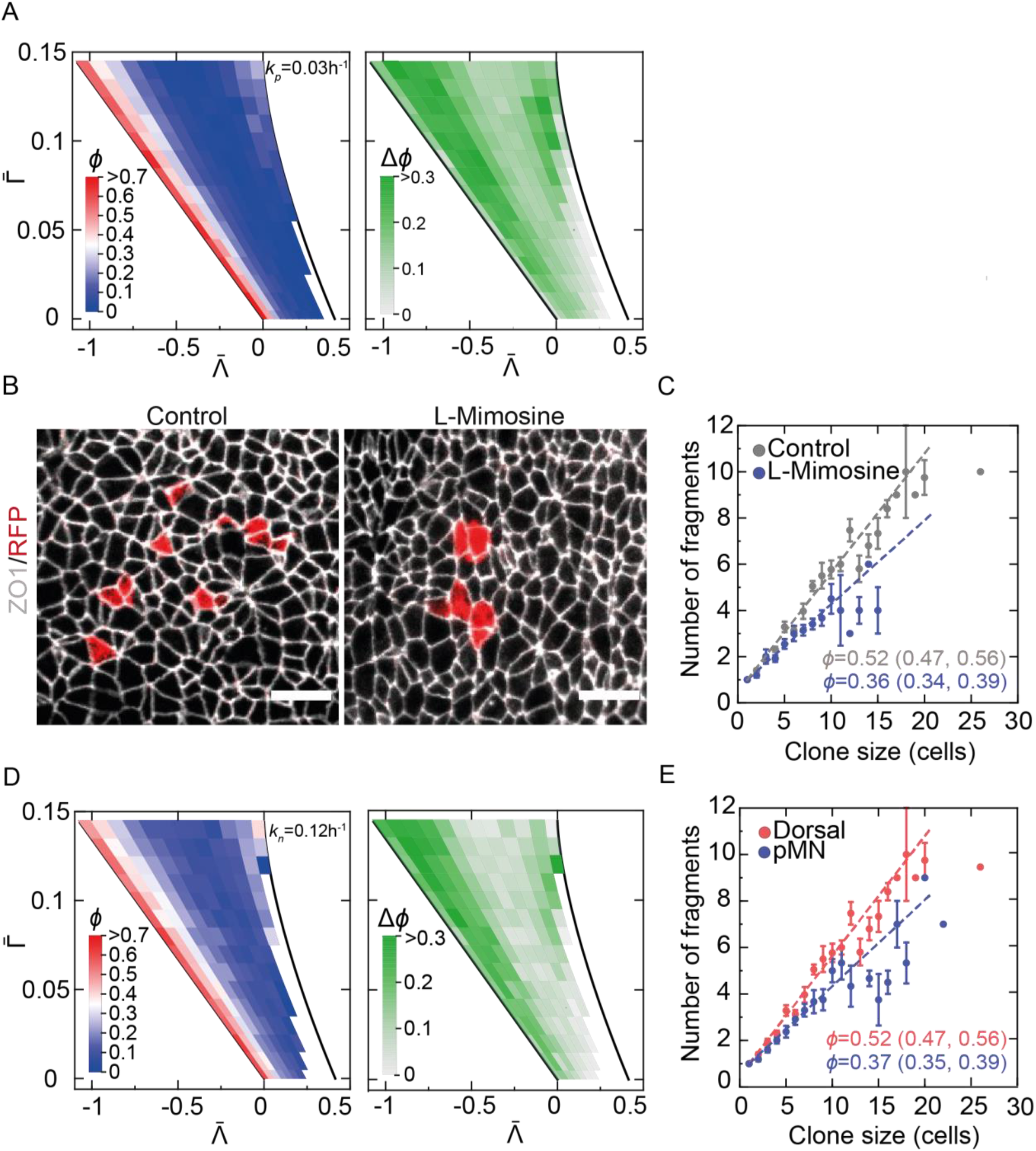
Clone fragmentation is determined by the proliferation and differentiation rates. **A**. Left, fragmentation coefficient for simulations with low proliferation rate *k*_*p*_ = 0.03h^-1^. Right, difference in *ϕ* between high (0.09h^-1^, Figure 2A) and low (0.03h^-1^) *k*_*p*_. **B**. Confetti clones induced at E7.5, embryos cultured from E8.5 for 42h with vehicle or 210 μM L-mimosine. ZO1 immunostaining, white. Scale bars, 10 μm. **C**. Mean number of fragments per clone for a given clone size and fragmentation coefficient (95% CI). **D**. Left, *ϕ* for simulations with high differentiation rate *k*_*n*_ = 0.12h^-1^. *k*_*p*_ = 0.09h^-1^. Right, difference in *ϕ* between *k*_*n*_ = 0 (Figure 2A) and *k*_*n*_ = 0.12h^-1^. **E**. Confetti clones generated in B in control condition. Fragments per clone ± SEM were quantified in the dorsal and pMN domains. *ϕ* (95% CI). Sample sizes, see Table S1.

This analysis predicts that artificially lowering the proliferation rate will lead to decreased cell rearrangements. To test this, we induced clones in Sox2CreERT2::Rosa26-Confetti mouse embryos. Tamoxifen injection at E7.5, when Sox2 levels are low, and the stochastic activation of one out of four possible Confetti reporters allowed us to achieve sparse clonal labelling and at the same time obtain a higher absolute number of clones per embryo than with the MADM system (see Methods). 24h after clonal induction we cultured the embryos ex vivo in the presence of the cell cycle inhibitors L-mimosine or aphidicolin for 42h and subsequently flat mounted them for analysis at E10.0 (Figures 4B, S8A). As expected, these treatments resulted in reduced mean clone sizes compared to control embryos (Figure S8B). Crucially, comparison of inhibitor to vehicle treated control embryos showed that for a given clone size, the number of fragments per clone was significantly reduced in both the L-mimosine and aphidicolin treated conditions (Figures 4C, S8C). These results are in agreement with the model prediction and confirm that the proliferation rate has a profound influence on the extent of cell rearrangements in the neuroepithelium.

Alongside with the decline in proliferation rate that occurs across the neural tube, neural progenitors begin to terminally differentiate and delaminate from the epithelium at E9.5 of development with rates that vary between the progenitor domains. In particular, the pMN domain differentiates at a higher rate than the other domains from E9.5 to E10.5 (*23*). To test the effect of terminal differentiation on cell rearrangements, we modelled cell loss in silico by randomly assigning a fraction of cells with a zero target area (see Methods). These simulations showed that cell loss leads to a decrease in clone fragmentation throughout the 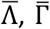 parameter space (Figure 4D).

To test whether this is consistent with experimental observations, we compared the fragmentation coefficients of Confetti clones at E10.5, 64h after tamoxifen injection, in the pD vs pMN domains. Consistent with the model predictions, clones in the pMN domain have significantly lower fragmentation coefficient (Figure 4E). Altogether, these observations indicate that the decline in cell rearrangements in the neural epithelium results from the temporal dynamics of the rate of proliferation and differentiation.

### Neuroepithelial cells maintain their tension and contractility over time

Our analysis showed that the decline in tissue growth rate has a strong effect on the dynamics of epithelial rearrangements. Nevertheless, it is possible that mechanical factors, such as tissue tension and/or contractility, could additionally contribute to reducing cell rearrangements during development. To test this, we analysed the cell shapes in ZO1 immunostained neural tubes from E8.5 to E11.5 (Figure 5A, see Table S3). We then compared them to simulations in order to infer 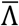 and 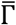, similar to previous studies (*30, 31*).

**Figure 5.**
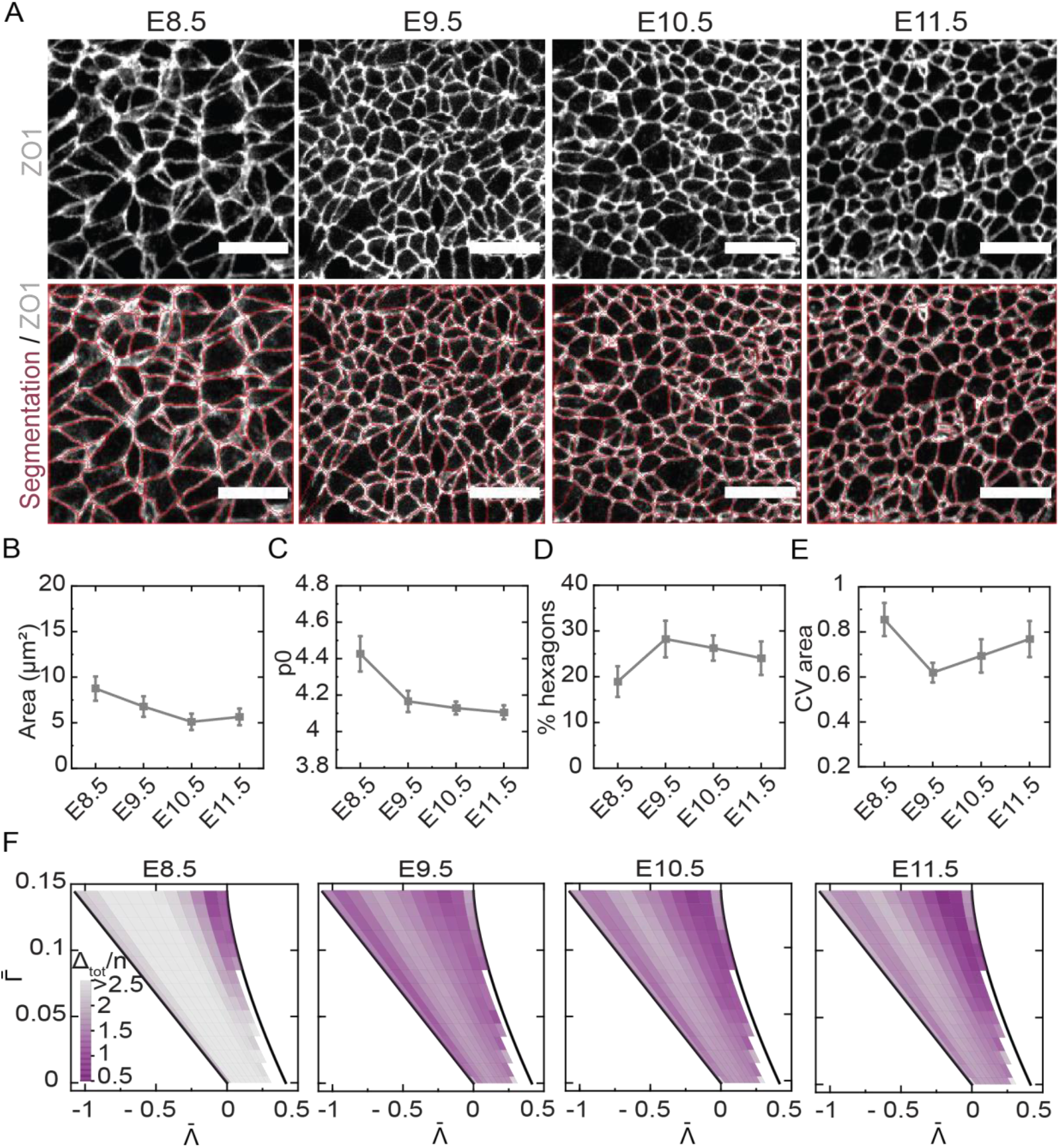
Cell shapes at different stages of neural tube development. **A**. Top, apical view of the epithelium with ZO1 immunostaining. Bottom, cell segmentation. Scale bars, 10 μm. **B-E**. Quantification of cell shape descriptors at different stages. **F**. Difference between the cumulative distribution of cell shape descriptors (p0, *α,hex*, 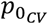, *ε*_*CV*_, *A*_*CV*_, *P*_*CV*_) between simulations and experimental data. Sample sizes, see Table S1.

Our data revealed that cell shapes change over developmental time. Consistent with increasing pseudostratification and the 2-fold increase in AB length of the epithelium from E8.5 to E10.5 (*23*), the apical cell area and perimeter decrease approximately 2-fold, maintaining constant cell volume (Figures 5B, S9A). Furthermore, p0 and cell elongation decline, while the fraction of hexagons increases (Figures 5C-D, S9B), indicating that cells become less elongated and more ordered over development. The area ratio slope, and the coefficients of variation of cell area, perimeter, and elongation also slightly decline (Figure 5E, S9C-E), but crucially, these parameters maintain overall high values throughout development that are closer to those observed in regions B and C, and least similar to region A (Figures 3C, S5). To compare directly the in vivo and in silico cell shapes, we plotted the cumulative difference in fraction of hexagons, area slope ratio, p0, p0 CV, elongation, elongation CV, area CV, perimeter CV between data and simulations across the 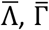 parameter space (Figure 5F, see Methods). This analysis confirmed that the cell shapes measured at E8.5 map to region C of parameter space. At E9.5, E10.5 and E11.5, the data is in good agreement with the simulations for a broader range of parameters, however the highest similarity is still in region C. Altogether, this analysis is consistent with the idea that that the neuroepithelium is characterized by high line tension and contractility, typical of region C, throughout developmental time.

To further test whether the tissue tension changes during developmental time, we performed laser ablation experiments at E8.5 and E11.5 of development using embryos expressing membrane localized fluorescent reporter in the neural epithelium (Figure S10A-B). There was no significant difference in the initial recoil velocity of the tissue between the two stages, suggesting that the active tension at these stages is similar (Figure S10C-D). Altogether, these results suggest that the dynamics of proliferation and differentiation, rather than changes in 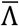 and 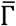, are the major factors that underlie the change in tissue fluidity over time.

Given that the neuroepithelium maps to region C of 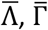 parameter space, we asked what is the significance of these mechanical characteristics for neural tube development. We therefore cultured E8.5 mouse embryos expressing Sox2-CreERT2::Rosa26:Confetti in vitro for 6 hours in 8μM ROCK inhibitor Y-27632, which impairs actomyosin organization and reduces tissue tension in the mouse neural tube (*32, 33*). Consistent with previous reports, ROCK inhibition impaired axis elongation and lead to enlarged cell areas and decreased apical F-actin accumulation within the neuroepithelium immediately after the ROCK treatment (Figure S10E-F), without impairing the ability of cells to divide. In order to track clones, we subsequently cultured the embryos without inhibitor for 36h and analysed clone fragmentation (Figure S10G-H). These data showed that the fragmentation coefficient of treated embryos (0.49 (0.38, 0.6)) was only marginally smaller than controls (0.55 (0.50, 0.61)), (Figure S10H). This suggests that a transient decrease in tissue tension was not sufficient to induce significant changes in cell rearrangements, reinforcing our findings on the specific and crucial role of cell proliferation in regulating tissue fluidity over time.

### The decline in cell rearrangements helps to maintain boundary precision

Our analysis indicates that the cell rearrangement rate declines after E9.5 of development. Previous work has shown that after E9.5, neural progenitor identities have been specified and are maintained by the underlying transcriptional network without requiring morphogen signalling (*23, 34, 35*). This raises the question what is the effect of growth on the precision of gene expression boundaries at these late stages.

To investigate this, we simulated a tissue that grows to a certain initial size, then marked two domains separated by a straight boundary. We then allowed the tissue to grow either with high proliferation rate (0.09 h^-1^) and hence high rate of cell rearrangements, or low proliferation (0.03 h^-1^), hence low rearrangement rate (Figure 6A). We measured the resulting boundary imprecision by its deviation from a straight line (here termed “wiggliness”) or the number of cells that become separated from the correct domain (“islands”, see Methods). Both measurements showed that after 16h the boundary imprecision was higher in the case of high cell rearrangements rate. In region C, the boundary wiggliness for high rearrangements rate corresponded to 3.43±0.09, compared to 1.94±0.05 for low rearrangements rate (Figure 6B). Similarly, the number of islands along a 50μm section of the boundary corresponded to 2.21±0.25 and 0.43±0.11 for high and low rearrangement rate, respectively (Figure 6C). These results suggests that the decline in cell rearrangements at later developmental stages may help to achieve the observed boundary precision in the neural tube (*34*).

**Figure 6.**
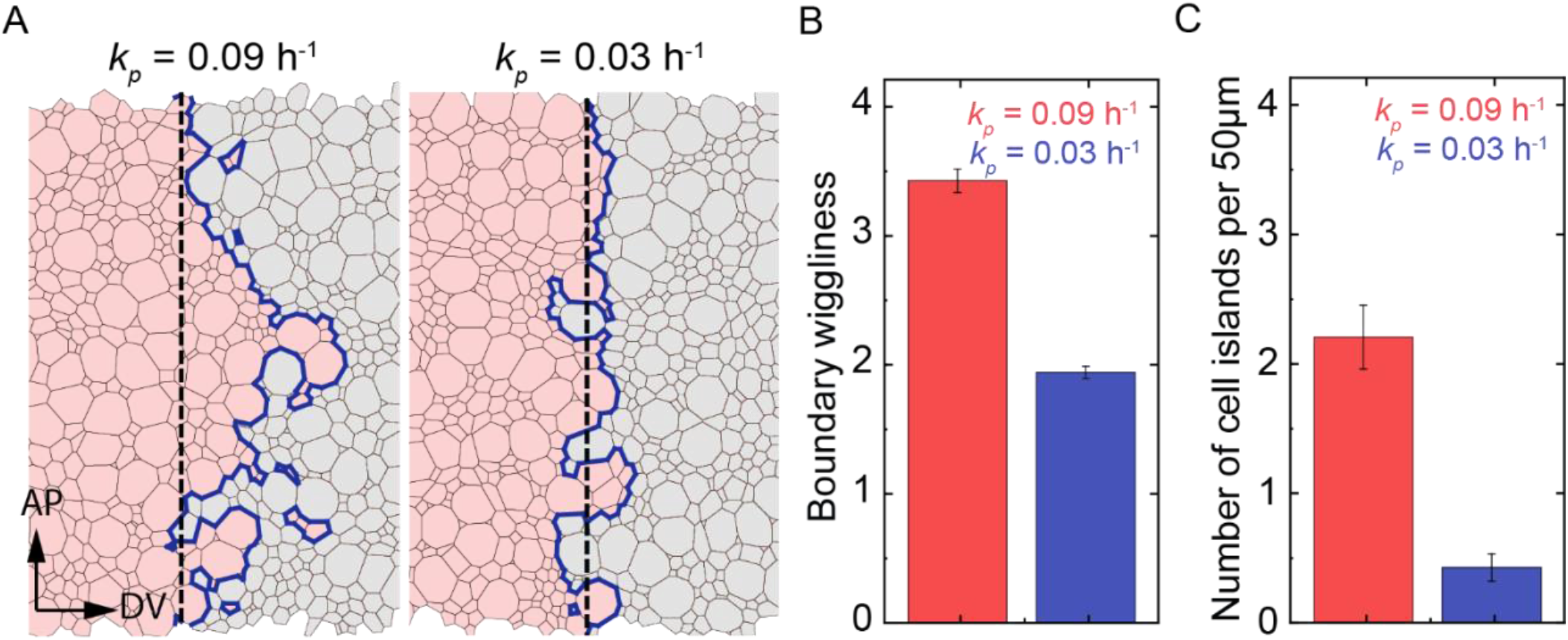
The rate of cell rearrangements influences boundary precision. **A**. Snapshots of domain boundaries in region 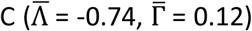 for high and low proliferation rate (*k*_*p*_ = 0.09h^-1^ and 0.03h^-1^, respectively) at the end of the simulations. Dashed line, *AP*_*length*_. Blue line, outer boundary of the red/gray domain. **B**. Boundary wiggliness after 16h simulation, region C. **C**. Number of islands per 50 μm in region C.

## Discussion

Morphogenetic processes have recently been linked to transitions in the material properties of tissues (*36*). Here we demonstrate that in the mouse neural tube epithelium, there is significant decline in tissue fluidity around E9.5 of development. Previous studies show that tissue rheology can change in the absence of noise or fluctuations (*3, 4, 37, 38*). In contrast, the transition we identified here resembles a glass transition, controlled by changes in active stresses within the tissue. We show that in the neuroepithelium, active stresses are generated by interkinetic nuclear movements during the cell cycle. Consequently, the proliferation rate determines the extent of cell rearrangements and tissue fluidity. The regulation of tissue fluidity has previously been linked to changes in cell density, mechanical properties, such as cell adhesion, cortical tension, contractility, as well as signaling pathways such as Wnt/PCP and Ffg (*2, 3, 5, 6, 37, 39–41*). By contrast, the role of cell divisions, apart from a few experimental examples (*3, 9, 10*) and theoretical predictions (*8, 42*), has been largely underappreciated. Given that cell cycle lengthening is observed in multiple developing tissues (*43–45*), our findings raise the possibility that these tissues also undergo effective cell-cycle driven increase in rigidity over time.

A specific feature of our analysis is the implementation of IKNM in the model. This revealed a novel fluid-like region, in which the tissue can proliferate and rearrange extensively, while at the same time maintaining the high tissue tension necessary for normal neural tube closure in amniotes (*32, 33*). The extent of cell rearrangements in mouse at early stages, albeit high (average clone spread of ∼3 cell diameters in 16h), is lower compared to cell movements that occur during secondary neurulation in zebrafish (one cell diameter per 10-20 minutes (*13*)). Furthermore, cell rearrangements at early stages do not lead to increased boundary imprecision when compared to the precision of morphogen gradients at these stages (*34*). Nevertheless, our simulations indicate that in the absence of morphogen signalling, the rate of cell rearrangements affects the straightness of domain boundaries. This suggests that the decreased rate of cell rearrangements at late stages may help to maintain high boundary precision. Altogether, these data reveal species-specific differences and raise the possibility that the primary neurulation observed in amniotes could be an adaptation that allows for precise pattern formation.

IKNM are characteristic of many epithelia (*46–48*), hence our conclusions might be relevant to other tissues besides the neural tube. Our analysis indicated that IKNMs exert their effect on cell rearrangements by affecting the apical surface area kinetics of cells. IKNM causes a rapid increase in the apical area just prior to division, dynamically changing the distribution of cell areas in the tissue which results in large cell area variation. Interestingly, in the Drosophila wing disc, the presence of smaller than normal mutant cells has been shown to induce clonal fragmentation (*24*). Yet, in our analysis, imposing “static” cell area variation did not lead to high clonal fragmentation, indicating a distinct mechanism whereby the kinetics of cell area changes are crucial for epithelial rearrangements.

## Methods

### Experiments

#### Mouse strains and generation of clones

The following strains were previously described: MADM-11^*TG*^ and MADM-11^*GT*^ (*22*), Rosa26-Confetti (Brainbow 2.1, (*49*)), Sox2-CreERT2 (*50*), mTmG (*51*). To generate MADM clones in MADM ^*TG/GT*^ trans-heterozygous Sox2-CreERT2 expressing embryos, MADM-11^*TG/TG*^ mice were bred to MADM-11^*GT/GT*^, Sox2-CreERT2/+ and pregnant females were injected with 3mg/mouse of tamoxifen. To generate Confetti clones, heterozygous Sox2-CreERT2 mice were bred to heterozygous Rosa26-Confetti and pregnant mothers were injected with 0.75mg/mouse of tamoxifen. Tamoxifen stock was prepared fresh in sunflower oil.

The first time point where we observe labelled cells is 8h after tamoxifen injection (Figure S1B), reflecting the time it takes for the nuclear translocation of Cre and subsequent onset of reporter expression. Thus, the time of Cre activity in the 24h tracing experiments (Figure 1) is considered 16h.

In the case of MADM clones, G2 recombination followed by X-segregation of chromosomes in mitosis produces a GFP and an RFP expressing daughter cell (Figure 1A). G2-Z segregation produces an unlabeled and a GFP/RFP coexpressing daughter cell. G1 recombination produces a GFP/RFP coexpressing cell. In our dataset, we found an increasing proportion of GFP/RFP coexpressing cells over time (Figure S1C), which correlates with the increasing relative G1 duration over time (*23*). This suggests that the majority of GFP/RFP clones result from G1 recombination.

#### Immunohistochemistry and imaging

For E9.5 and later stages, embryos were bisected along the roof plate prior to fixation and along the floor plate prior to staining. Embryos were fixed, immunostained and the brachial region flat mounted as previously described (*23*), with 10h washes in PSB, 0.1% Tween after primary and secondary antibody incubations. Primary antibodies used were: mouse anti-ZO1 (Invitrogen, 1:90), goat anti-Olig2 (R&D systems, 1:100), sheep anti-GFP (AbD Serotec, 1:1000), rabbit anti-RFP (Rockland, 1:2000), mouse anti-Nkx2.2 (DSHB, 1:20). Secondary antibodies: donkey anti-mouse Alexa fluor 647 and donkey anti-goat FITC (Jackson Immuno, 1:250), donkey anti-rabbit Cy3 (Jackson Immuno, 1:1000).

For MADM clone analysis, embryos were immunostained against ZO1, RFP, GFP and Nkx2.2. Clones located within a 25μm interval dorsal to the Nkx2.2 domain were considered to be pMN clones. For Confetti clone analysis, embryos were immunostained against ZO1 and Olig2. To stain actin filaments, the following steps of the protocol were modified: embryos were fixed in 4% PFA overnight, methanol fixation was omitted, and Alexa fluor™ 488 Phalloidin (ThermoFisher Scientific, 1:100) was added together with the secondary antibody.

Imaging was performed using a 40X/1.3 NA oil objective on a LSM880 inverted confocal microscope. Images of the apical surface capturing the entire dorsoventral length of the epithelium were acquired through tile scanning with Z-slices 0.8μm apart. Tiles were configured in the form of a grid and overlapped 10%. Subsequently, the tiles were stitched using the BigStitcher plugin in Fiji.

#### Mouse embryo culture and inhibitor treatments

Heterozygous Sox2-CreERT2 mice were bred to Rosa26-Confetti and pregnant mothers were injected with 0.75mg/mouse of tamoxifen at E7.5 to induce clonal labelling. Sox2-CreERT2 induction at E7.5, when Sox2 levels are low, and the stochastic activation of one out of four possible Confetti fluorescent reporters allows us to achieve sparse clonal labelling but obtain a higher absolute number of clones per embryo than with the MADM system. 24h later, at E8.5, embryos were dissected and cultured with their yolk sac intact in temperature controlled roller culture as previously described (*52*). In order to perturb proliferation, embryos were cultured in the presence of 210μM L-mimosine (Sigma) or 800nM Aphidicolin (Sigma) for 42 hours. For ROCK inhibition, embryos were cultured in the presence of 8μM ROCK inhibitor Y-27632 (Calbiochem) for 6h, subsequently the medium was replaced to default medium without ROCK inhibitor and the embryos were cultured for another 36h. After culture, embryos were harvested and processed for imaging as described above.

#### Laser ablation

Homozygous mTmG mice were crossed Sox2-CreERT2 and pregnant mice were injected with 3mg Tamoxifen 24h prior to harvesting the embryos. Upon recombination, membrane GFP was specifically expressed in the neural epithelium, and this was used to localize the tissue. Laser ablation measurements were performed using the tdTomato signal, which had a better signal to noise ratio than GFP. To perform the laser ablation, whole E8.5 embryos and bisected and flat mounted E11.5 neural tubes were immobilized for live imaging using fibrinogen-thrombin clots as previously described (*23*). The samples were kept in whole embryo culture medium in an environmental chamber with 5% CO2 at 37C. Laser ablation was performed on an Andor spinning disc system with inverted Axio Observer Z1, C-Apochromat 40x/1.2 water immersion objective (Carl Zeiss) using a 355 nm pulsed UV-A nano-laser (Teem photonics) at 1.6-1.8% laser power with 25 pulses (2 shots/μm) at 1000 Hz. The tissue was cut along a 25μm line oriented along the AP axis at intermediate dorsoventral positions. Images were collected with 250ms exposure time and frame rate of 1.5s. For determining the recoil velocities, the ablated edges were identified from the fluorescence intensity peaks measured using the Fiji plot profile function, and tracked over time. The displacement of the edges was measured with respect to the line of the cut at t=0. The initial recoil velocity was computed by dividing the initial displacement by the duration of the first frame (1.5s).

### Data analysis

#### Clone identification procedure

Images were processed in Fiji. Labelled progenitor cells were manually marked at their apical surface at the level of the ZO1 staining. Fragments were defined as groups of adjacent cells that share an edge or a vertex. Clones were defined as groups of labelled progenitor cells in close proximity of each other (<25μm to the nearest labelled cell). Images of MADM clones were processed manually. In the case of Confetti clones, only the RFP, YFP and CFP reporters, which can be detected at the apical surface, were used for analysis, while clones labelled by the nuclear GFP were excluded. The sparseness of labelling in the experiments was as follows: 108±9, 136±10 and 81±9 cells/mm^2^ (equivalent to 16±1, 20±1, and 12±1 clones/mm^2^) for CFP, RFP, and YFP, respectively (mean±SEM for 155 images is given). To identify Confetti clones, cell coordinates were recorded and subsequently analysed using a custom-built python script. CFP, RFP and YFP channels were analysed separately. Labelled cells were assigned to the same fragment if the distance between them was <5 μm and to the same clone if they were <25μm apart. These assignments were consistent with visual identification of fragments and clones. Labelled postmitotic neurons that have delaminated from the neural epithelium were excluded from our analysis.

#### Fragmentation coefficient estimation

We determined the fragmentation coefficient *ϕ* by fitting *f* = *ϕs* + *b* to the respective dataset, where *f* is the mean number of fragments for a given clone size, *s* is the size of the clone in cells, and *b* is an offset parameter that is chosen in such a way that the line crosses through the point (1,1), reflecting the fact that single cell clones have one fragment by definition. For clones analysed 24h after tamoxifen injection, reliable statistics could be obtained for clones with 4 cells or less, hence only these clone sizes were used for estimating *ϕ*. In mouse embryo culture experiments, clones were analysed 64h after tamoxifen injection. In this case, reliable statistics could be obtained for clone sizes up to 8 cells and these were used to estimate *ϕ*.

#### Growth rate estimation

The growth rate of MADM clones *k*_*g*_ was inferred from the mean clone size *s* as *k*_*g*_ = In(*s*) /Δ*t*, where Δ*t* = 16h is the time interval of Cre activity in the experiments (Figure S1B and Methods).

#### Spread and anisotropy of clones

To estimate the spread of clones, the coordinates of cell centers in a clone were used to determine the clone centroid. The maximum spread of the clone was quantified as the distance between the clone centroid and the furthest cell center. To estimate the mean maximum spread for a given developmental stage, clones of all sizes were taken into account (including one cell clones).

Clone anisotropy was quantified by drawing a bounding rectangle around the clone, using the apical cell outlines, marked by ZO1, to demarcate cells. Images are always oriented so that the vertical axis is aligned with the tissue DV axis. The aspect ratio of the clone is then given by the DV to AP side lengths of the bounding rectangle. Note that quantifying the clone shape at the apical surface, rather than the cell bodies or nuclei, avoids potential artefacts of tissue mounting, whereby clone shape could be affected by misalignment of the apical and basal surfaces of the neural epithelium.

#### Segmentation of cell shapes

The apical surface of E8.5, E9.5, E10.5 and E11.5 embryos was segmented using the Tissue analyzer (TA) plugin (*53*) in Fiji. Cell outlines were automatically identified and manually checked for correctness. TA plugin provided description of polygonal mesh including vertices and edges of cell outlines as well as the number and identify of cell neighbours. The cell area was calculated using standard formula for the area of the n-gon and cell perimeter as sum of length of polygon edges. Cell elongation was calculated as in (*26, 30*).

### Simulations

#### Vertex model description and implementation

The vertex model used in this study has previously been described (*26*) and was implemented here using Python 3.7. Briefly, the following energy function is minimized in every simulation step:

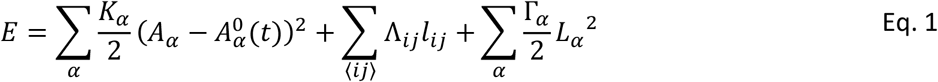

where *α* = 1, …, *N*_*c*_ enumerates all cells, *i* = 1, …, *N*_*v*_ enumerates all vertices, *K*_*α*_ is elasticity coefficient, *A*_*α*_ is area of cell *α*, and 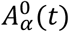 is the preferred area of cell *α* at time *t*, Λ_*ij*_ is the line tension coefficient associated with cell edge between *i* and *j* of length *l*_*ij*_, and Γ_*α*_ is the contractility coefficient of cell *α* with perimeter *L*_*α*_. We assume that parameters are the same for each cell (*K*_*α*_=*K, Γ*_*α*_=*Γ*), and for each edge (Λ_*ij*_=Λ) if no noise in the line tension is considered. 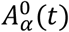 is a piecewise linear function reflecting the effect of IKNM on the apical cell area in the 4 phases of the cell cycle (G1, S, G2, M, also see below). Adopting the same notation as previous studies (*26, 27*), we use normalized parameters 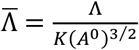 and 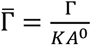 where *A*^0^ is the average target area during the cell cycle.

The motion of vertices is determined from the first-order kinetics: 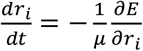, where *r*_*i*_ is position of vertex *i*, and *μ* is the drag coefficient. Tissue growth was considered to be anisotropic with drag coefficients *μ*′ and *μ*′′ in DV and AP directions, respectively (*26*).

The following changes were made in the current version of the model:

##### Implementation of junctional noise

We considered that fluctuations in the internal line tension follow an Ornstein-Uhlenbeck process, 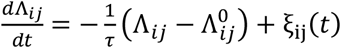, where *ξ*_*ij*_ (*t*) is white, uncorrelated noise with ⟨*ξ*_*ij*_ (*t*) ⟩ = 0 and 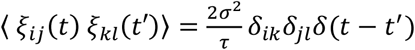. We used the following discretization (*29*):

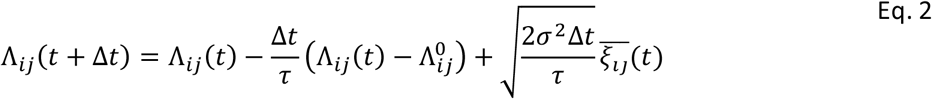

where Δ*t* is a time step used in the simulation, *τ* is the line tension correlation time, *σ* is the intrinsic line tensions deviation, 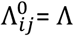 is a reference line tension that corresponds to the line tension without noise, and 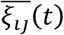 is drawn from the Gaussian distribution *N*(0,1).

##### Implementation of T1 and T2 transitions

The T1 transition is defined as in (*26*). In particular, when an edge between two neighbouring cells is shorter than a predefined small length *l*_*T*1_, this edge is replaced with a new edge that is perpendicular to the old edge and has a length *l*_*new*_ = 1.01 *l*_*T*1_. Using this definition, we observed that for negative line tension or in the presence of line tension fluctuations, immediately after a T1 transition the new edge can shrink instead of extending, thus leading to a reverted T1 transition. This can occur multiple times at a given edge, hence we call this an oscillatory T1 transition.

One strategy to partly mitigate the occurrence of oscillatory T1 transitions is to increase *l*_*new*_/*l*_*T*1_ ratio (*54*). However, particularly in region A, the oscillatory T1 transitions are generic and increasing *l*_*new*_/*l*_*T*1_ does not result in decreasing the number of oscillatory T1 transitions. Therefore, instead we approached this by keeping track of the oscillatory T1 transitions and subtracting them from the overall count of T1 events in our statistics. More specifically, we track how many T1 transitions occurred for every edge, using the dictionary data structure in Python. If repeated T1 transitions occurred *n*_*T*1_ times between time *t*_0_ and *t*_*nT*1_, their contribution to the T1 unique rate, T1_UNQ,_ was considered to be 1/*n*_*T*1_ for times between *t*_0_ and *t*_*nT*1_.

T2 transitions are defined as in (*26*). In particular, cells in which the area becomes very small, have shrinking edges. This results in sequential T1 transitions, which finally lead to a double-sided cell with zero area. Such cells are removed from the simulation by merging the two vertices that delimit the double-sided cell into one vertex. The last T1 transition that results in double-sided cell is counted as a T2 transition, and is not included in overall number of T1 transition events. All T1, T1_UNQ_ and T2 rates are estimated in time windows of Δ*t*=2h by dividing number of respective events by the average number of cells in this time window. The T1 rate reported in the main text and in Figures 2B, 3B and S7B is defined as T1 unique rate.

The cell removal from the tissue through differentiation is implemented similar to (*26*) with the additional requirement that if cell was randomly selected for differentiation, the line tension coefficients Λ_*ij*_ for this cell are no longer fluctuating and have assigned positive value Λ_*ij*_=0.2 which fosters shortening of all edges of this cell.

##### Cell lineage tracing

In order to efficiently analyse in silico clonal populations, the complete information about cell lineage, i.e. daughter cell identifiers and division times, are stored. For the analysis of clone fragmentation in silico, we used all clones per simulation and 10 independent simulations per parameter set.

##### Parameters of the model

The parameters used are summarized in Table S2. The specific proliferation (*k*_*p*_) and differentiation (*k*_*n*_) rates used are given in the figure legends. Every data point across the 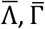 parameter space was obtained by pooling together cells from 10 independent simulations for a given set of parameters 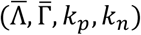.

##### Vertex model initialization and in silico clone tracing

Similar to (*26*), the vertex model is initiated with a regular hexagonal lattice of 10 rows with 10 cells/row. In the initial simulation phase, the tissue grows for 16h with 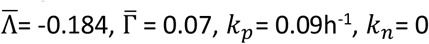, and *σ* = 0. After the initial simulation phase, the number of cells is 460 ±19 (mean ±SE), the time is set to 0, and parameters are modified to target simulation parameters. The tissue is then allowed to grow with the target simulation parameters for 8h, subsequently clones are labelled and clonal populations of cells are tracked for 16h.

#### Analysis of the contribution of cell area heterogeneity and kinetics to clonal fragmentation

To investigate the area variability of cells undergoing a T1 transition, we defined quadruplets of neighbouring cells, designated A, B, C, D, where A and B share a common edge, and C and D do not (see schematic in Figure. 3D). We further define the cell names based on the cell area, so that the area of A < area of B and area of C < area of D. If the common edge between A and B shrinks below *l*_*T*1_, a T1 transition takes place, as a result of which A and B are no longer adjacent, whereas C and D become new neighbours. For comparison, ‘random’ quadruplets are generated by randomly finding A and B cells separated by a common edge, and finding cells C and D that are adjacent to A and B, but not to each other. Note that because the polygonal mesh has no rosettes, that is each vertex has 3 edges associated with it, the assignment of a quadruplet to an edge is unique.

##### Cell area kinetics during the cell cycle and cell division

The interkinetic nuclear movement (IKNM) is approximated as a linear combination of two terms, one corresponding to linear increase in cell volume and the other interpolating for the change in apical cell surface as a function of age of a cell,

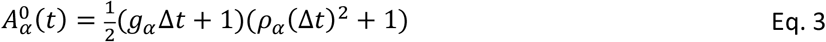

where *g*_*α*_ is the growth rate of the cell *α*, Δ*t* = *t* − *t*_*new*_ is age of the cell that divided at *t*_*new*_, and *ρ*_*α*_(Δ*t*) is piece-wise linear function representing the apical-basal position of the nucleus as a function of Δ*t* and cell cycle phase (*26*). This function equals zero in S-phase of the cell cycle, in which nucleus stays basal, and 1 during the mitosis, when the nucleus is apical. The exact form of *ρ*_*α*_(Δ*t*) is defined in (*26*). The growth rate *g*_*α*_ is drawn from a normal distribution with mean equal to 1/*t*_*T*_ where *t*_*T*_ is the total cell cycle time, and standard deviation *σ*_*g*_=0.45/*t*_*T*_. Negative growth rates are not allowed. In the simulation the proliferation is defined as *λ* = In 2/*t*_*T*_. The cell divides when cell is in the M-phase, 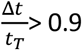, and cell volume exceeds a critical value, 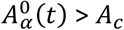. The cell divides by introducing a new edge that splits this cell into two daughter cells that enter the next cell cycle (*26*).

In order to study how the cell area kinetics affects T1 transitions, we use different forms of 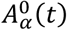. In Figure 3E-G different conditions for area kinetics are defined as in Table S4. All conditions except for ‘IKNM’ condition have noise in the line tension as in eq. 2 with *τ*=37s, *σ*=0.02.

#### Comparison between model and data

In order to compare results of vertex model simulations to experimental data, we use the cumulative distance between model and experimental data including the following nondimensional descriptors *D* ∈ {*p*_0_, *ϵ, α, hex, p*_0*CV*_, *ϵ*_*CV*_, *A*_*CV*_, *P*_*CV*_ } (see Table S3). For each descriptor *D*, we calculate a difference 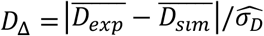, where 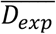 is the mean value of *D* obtained for all cells in the simulation for a given set of parameters 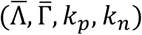 at final time,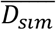 is the mean value of *D* estimated for the segmented data at a specific developmental stage, and 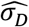 is standard deviation of 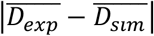 over all samples. The cumulative distance Δ_*tot*_/*n* is then defined as the sum of the differences *D*_Δ_ for all descriptors, normalized to the number of descriptors, i.e.: 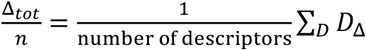.

#### Simulations and measurement of boundary precision

Two types of cells arranged in two domains separated by a straight line at *x* = 0 are marked at 8 h. The model has double periodic boundary conditions, hence the cells are uniquely identified within a reference box with dimensions *x* ∈ (−*DV*_*length*_, +*DV*_*length*_), and *y* ∈ (0, *AP*_*length*_). Thus, cells in one domain have *x* ∈ (−*DV*_*length*_, 0), all remaining cells are in the other domain at *t* = 8 h. The boundary shape is subsequently measured after 16h of tracking.

‘Islands’ are defined as connected components of the cell network for each cell type, excluding the largest component, which corresponds to the main domain. In order to compare the number of islands for tissues of different sizes, the number of islands is divided by 2*AP*_*length*_. The number islands for the two domains are independent and both are used to calculate the average number of islands.

‘Wiggliness’ is defined as the total length of the outer boundary of the domain divided by 2*AP*_*length*_ at the final time. For each domain, the outer boundary is a sum of the lengths of the two boundaries of the main domain, and the boundaries of islands of both cell types. In order to compare wiggliness for tissues of different sizes, the total length of edges constituting the outer boundary is divided by 2*AP*_*length*_. Note that the wiggliness of the two domains is the same.

## Acknowledgements

We thank S. Hippenmeyer for reagents, CP Heisenberg, J. Briscoe, and K. Page for comments on the manuscript. Work in the AK lab is supported by IST Austria, the European Research Council under European Union (EU) Horizon2020 research and innovation program grant 680037, and Austrian Science Fund (FWF): F78 (Stem Cell Modulation). MZ was supported by the Narodowe Centrum Nauki, Poland (SONATA, 2017/26/D/NZ2/00454) and the Polish National Agency for Academic Exchange.

## Author contributions

LB performed experiments, quantified and analysed data; AS performed and analysed laser ablation experiments; MZ performed computational analysis; LB, AS, EH, MZ, AK designed experiments; AK, EH, MZ analysed and interpreted results; AK conceived the project and wrote the manuscript.

## Additional tables and figures

**Table S1.** Sample sizes.

| Condition | Number of clones |
| --- | --- |
| MADM E8.5 +24 h | 46 |
| MADM E9.5 +24 h | 87 |
| MADM E10.5 +24 h | 94 |
| L- mimosine treatment | 155 |
| Control condition of L-mimosine treatment | 382 |
| Dorsal domain at E10.0 | 319 |
| pMN domain at E10.0 | 190 |

B. Number of segmented cells analyzed for Figure. 5A-B and figure. S9.
| Condition | Number of segmented cells |
| --- | --- |
| Apical surface of E8.5 neural epithelium | 6929 |
| Apical surface of E9.5 neural epithelium | 2433 |
| Apical surface of E10.5 neural epithelium | 5520 |
| Apical surface of E11.5 neural epithelium | 3583 |

**Table S2.** Vertex model parameters and their values in simulations.

| Parameter | Description | Value |
| --- | --- | --- |
| $\Delta t$ | Simulation time step | 0.29 s |
| $\Delta l$ | Simulation length unit | 4.6 $\mu\text{m}$ |
| $K$ | Elasticity | 1 au |
| $\bar{\Lambda}$ | Normalized line tension | $\bar{\Gamma} = 0, 0.01, \dots, 0.14$<br>for each $\bar{\Gamma}$ there are 10 equally spaced $\bar{\Lambda}$ that are delimited by:<br>$-4\sqrt{2}3^{1/4}\bar{\Gamma} < \bar{\Lambda} < \frac{2\sqrt{6}}{27}(\sqrt{3} - 12\bar{\Gamma})^{3/2}$ |
| $\bar{\Gamma}$ | Normalized contractility | |
| $A_c$ | Critical cell area | 27 $\mu\text{m}^2$ |
| $l_{T1}$ | T1 transition length threshold | 0.046 $\mu\text{m}$ |
| $l_{new}$ | Edge length after T1 transition | $1.01l_{T1}$ |
| $\mu$ | Medium viscosity | 0.12 $K \mu\text{m}^2 \text{s}$ |
| $\mu''/\mu'$ | AP to DV drag viscosity ratio | 50 |
| $t_T$ | total cell cycle time | 8 h, 12 h, 20 h |
| $k_p$ | proliferation rate, $\ln 2/t_T$ | 0.09 $\text{h}^{-1}$, 0.06 $\text{h}^{-1}$, 0.03 $\text{h}^{-1}$ |
| $k_n$ | terminal neuronal differentiation rate | 0, 0.12 $\text{h}^{-1}$ |
| $\sigma$ | magnitude of noise in internal line tension | 0.02 |
| $\tau$ | internal line tension correlation time | 37 s |
| $\sigma_g$ | standard deviation of cell growth rate (see eq. 3) | $0.45/t_T$ |

**Table S3.** Cell shape descriptors and their definition.

| Descriptor symbol | Name | Definition |
| --- | --- | --- |
| $A$ | Cell area | Apical surface area of the cells at the level of tight junctions ($\mu\text{m}^2$) |
| $P$ | Cell perimeter | Perimeter of the cells at the level of tight junctions ($\mu\text{m}$) |
| $p_0$ | Cell shape index | $p_0 = P/\sqrt{A}$ |
| $\varepsilon$ | Cell elongation | $\sqrt{v_1/v_2}$, where $v_1$ and $v_2$ are the eigenvalues of the second moment matrix of the vertices of the cell, $v_1 \geq v_2$ |
| $A_{CV}$ | Cell area CV | Coefficient of variation of $A$ |
| $P_{CV}$ | Cell perimeter CV | Coefficient of variation of $P$ |
| $p_{0\_CV}$ | Cell shape index CV | Coefficient of variation of $p_0$ |
| $\varepsilon_{CV}$ | Cell elongation CV | Coefficient of variation of $\varepsilon$ |
| $hex$ | Fraction of hexagons | Ratio of hexagons to all polygons in the tissue |
| $\alpha$ | Area ratio slope | Determined from a fit $\frac{\langle A_n \rangle}{\langle A \rangle} = \alpha(n - 3) + b$, where $\langle A_n \rangle$ is the mean area of n-sided cells and $n$ is the number of sides, $4 \leq n \leq 8$ |

**Table S4.** Simulations of cell area kinetics.

| Condition name | Description |
| --- | --- |
| IKNM +noise | $A_{\alpha}^0(t)$ as in eq. 3 |
| IKNM | $A_{\alpha}^0(t)$ as in eq. 3, no noise in the line tension, $\sigma=0$ |
| $\Lambda$ noise | $\rho_{\alpha}(\Delta t) = 1$ in eq. 3 for all stages in the cell cycle |
| $A_0$ noise | $\rho_{\alpha}(\Delta t) = z$ where $z$ is drawn from uniform distribution ranging from 0 to 2 |
| exp growth $\lambda$<br>exp growth $2\lambda$<br>exp growth $4\lambda$ | linear increase in cell volume is replaced with exponential increase, that is $A_{\alpha}^0(t) = \frac{1}{2}(\exp(g_{\alpha}\Delta t) + 1)$, with no IKNM; different conditions correspond to $g_{\alpha}=1, 2$, and $4$. |

**Figure S1.**
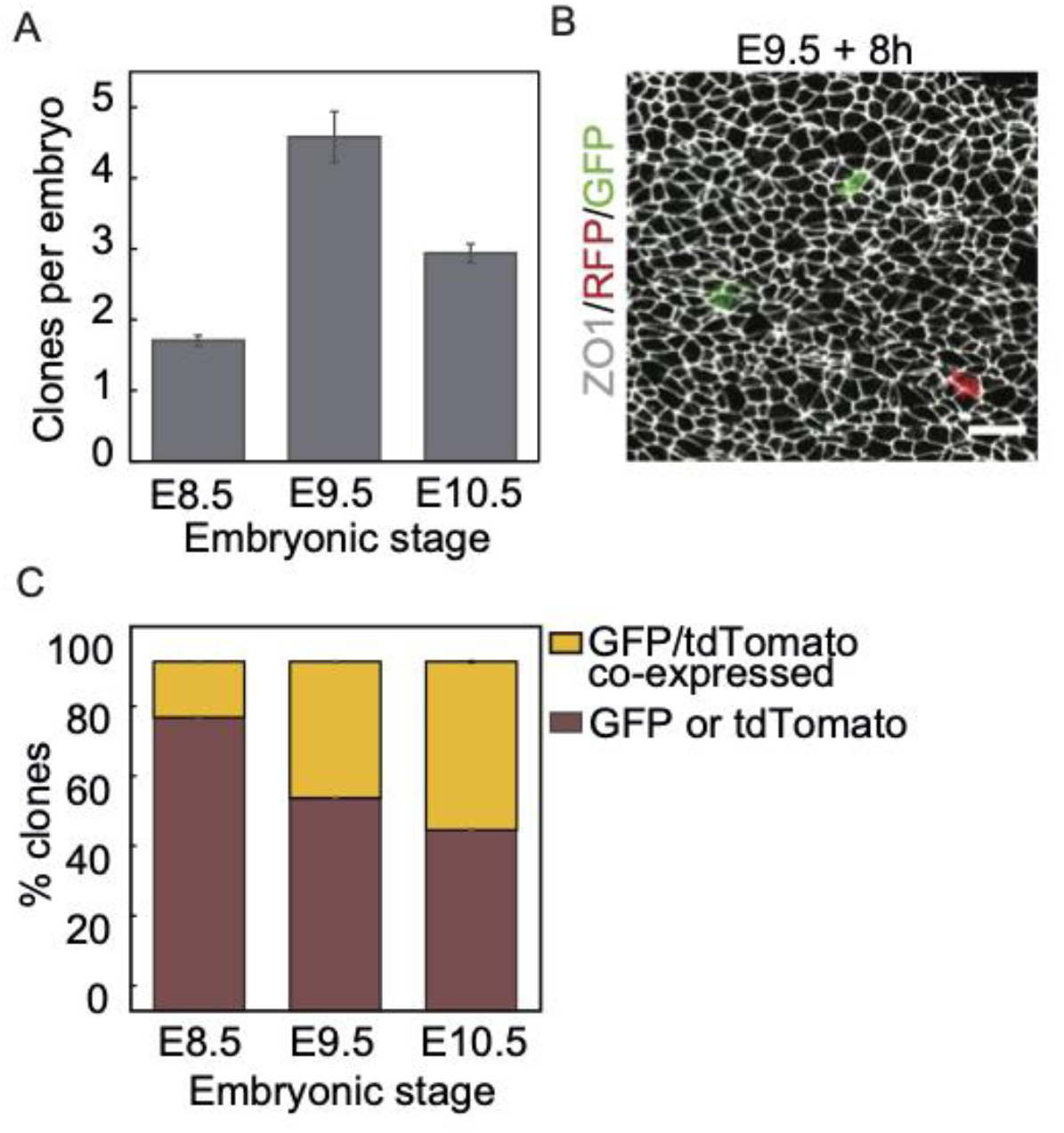
Clonal labelling in the mouse spinal cord. **A**. Mean number of MADM clones per embryo at different stages. **B**. Confetti clones generated at E9.5 after 8h of induction with tamoxifen. The tissue was immunostained against RFP, EGFP and ZO1. Scale bar, 10 μm. **C**. Fraction of MADM clones resulting from G2-X segregation, in which daughter cells expressed either GFP or tdTomato (brown), and clones with cells that co-expressed GFP and tdTomato resulting from G2-Z or G1 recombination (yellow). Error bars, mean ± SEM.

**Figure S2.**
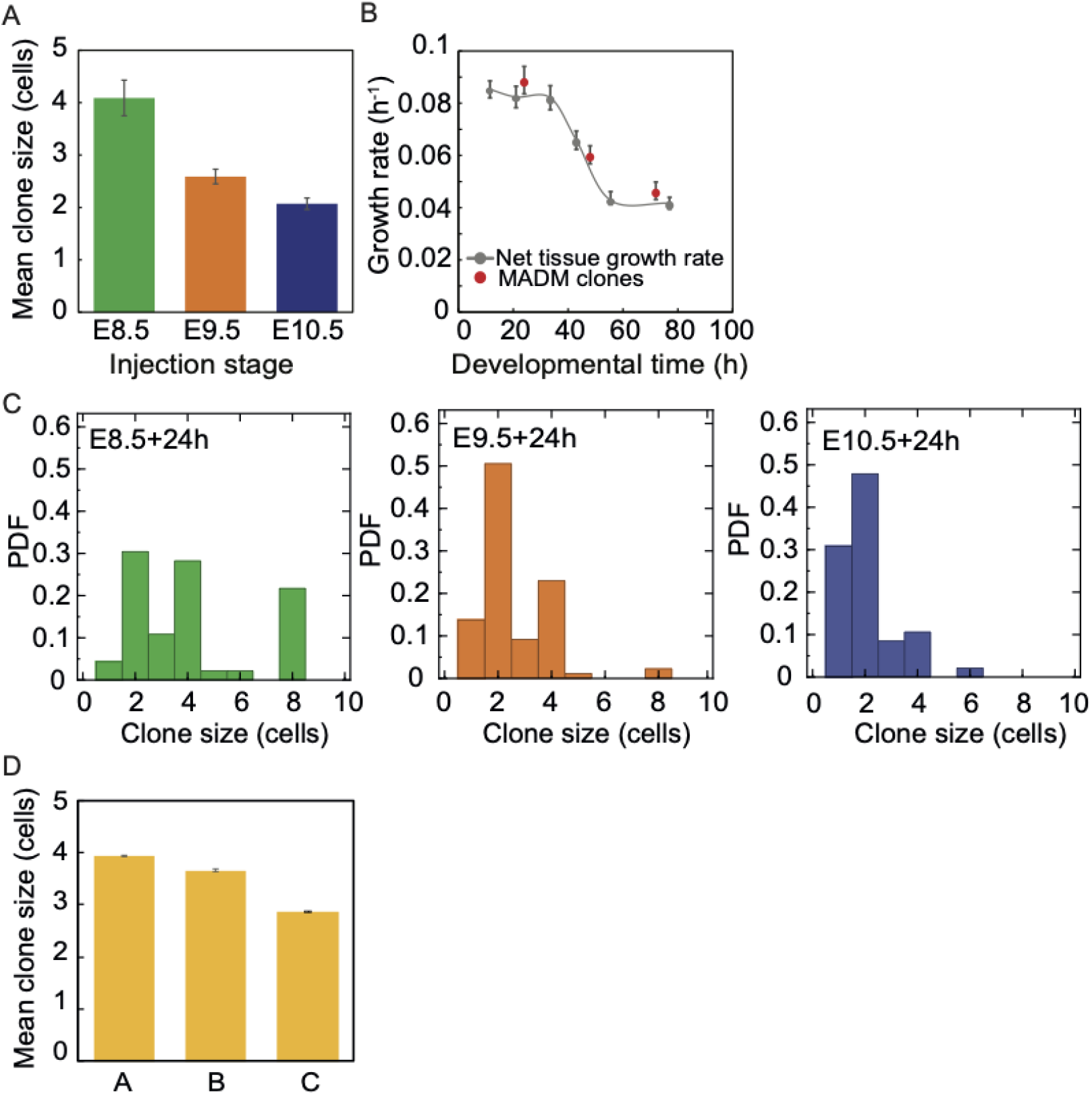
MADM clones reflect the tissue growth grate. **A**. Mean clone size generated at different stages. MADM clone size decreases over time. Error bars, mean ± SEM. **B**. Comparison of the growth rate measured from the net growth rate of the tissue in (*5*) and the growth rate predicted from the mean MADM clone size. The clonal growth rate *k*_*g*_ was estimated considering that clone size (*c*) grows exponentially, thus 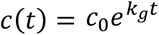. **C**. Clone size distributions at different stages. **D**. Mean clone sizes of in silico generated clones representative of regions A, B and C of parameter space. The clones were tracked for a period of 16h with proliferation rate *k*_*p*_ = 0.09 h^-1^. Simulation parameters were 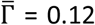, and 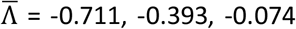 for regions A, B, C, respectively.

**Figure S3.**
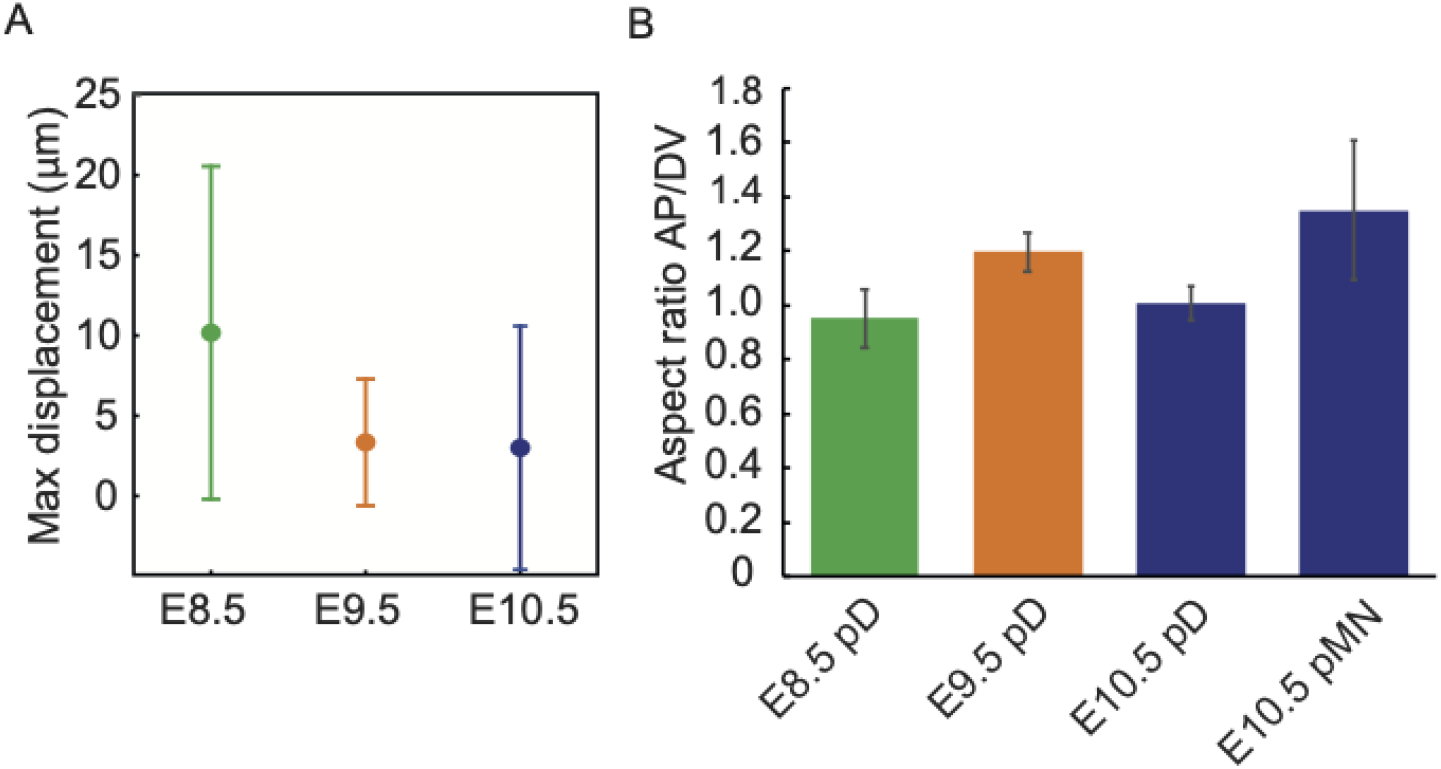
Spread of MADM clones. **A**. Maximum displacement of labelled cells from the clone centroid for clones induced at E8.5, E9.5 and E10.5. Maximum displacement at different stages correspond to 3.1±0.4 cell diameters, 1.1±0.1 cell diameters and 1.2±0.3 cell diameters, respectively. Mean±SEM. **B**. AP/DV aspect ratio of the clones at different developmental stages. Clones in the most ventral regions (p3 domain and floor plate) were excluded from the analysis.

**Figure S4.**
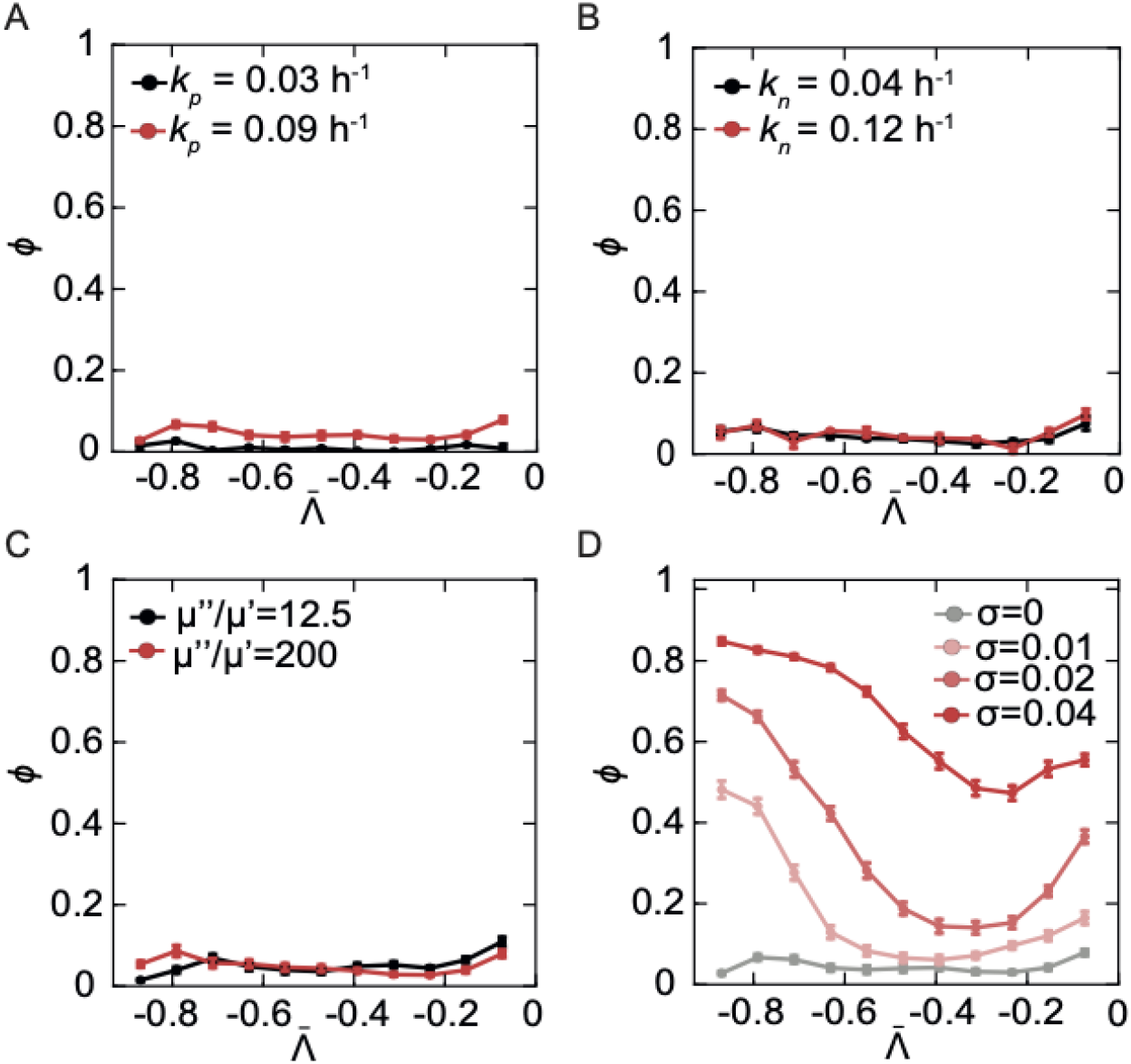
Clone fragmentation depends on line tension noise. **A-D**. Fragmentation coefficient *ϕ* in simulations performed for a fixed value of 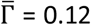 and different values of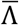. **A**. Variable proliferation rate *k*_*p*_, no line tension noise (*σ* = 0). **B**. Variable differentiation rate *k*_*n*_, no line tension noise (*σ* = 0). **C**. Variable ratio *μ*^′′^/*μ*^′^ of the drag viscosity in AP and DV directions, which changes the tissue growth anisotropy (*8*). *σ* = 0. **D**. The magnitude of noise in internal line tension is varied *σ* = 0, 0.01, 0.02, 0.04 with fixed internal line tension correlation time *τ* = 37s. The proliferation rate is *k*_*p*_ = 0.09h^-1^, no differentiation, and AP to DV drag viscosity ratio *μ*^′′^/*μ*′ = 50.

**Figure S5.**
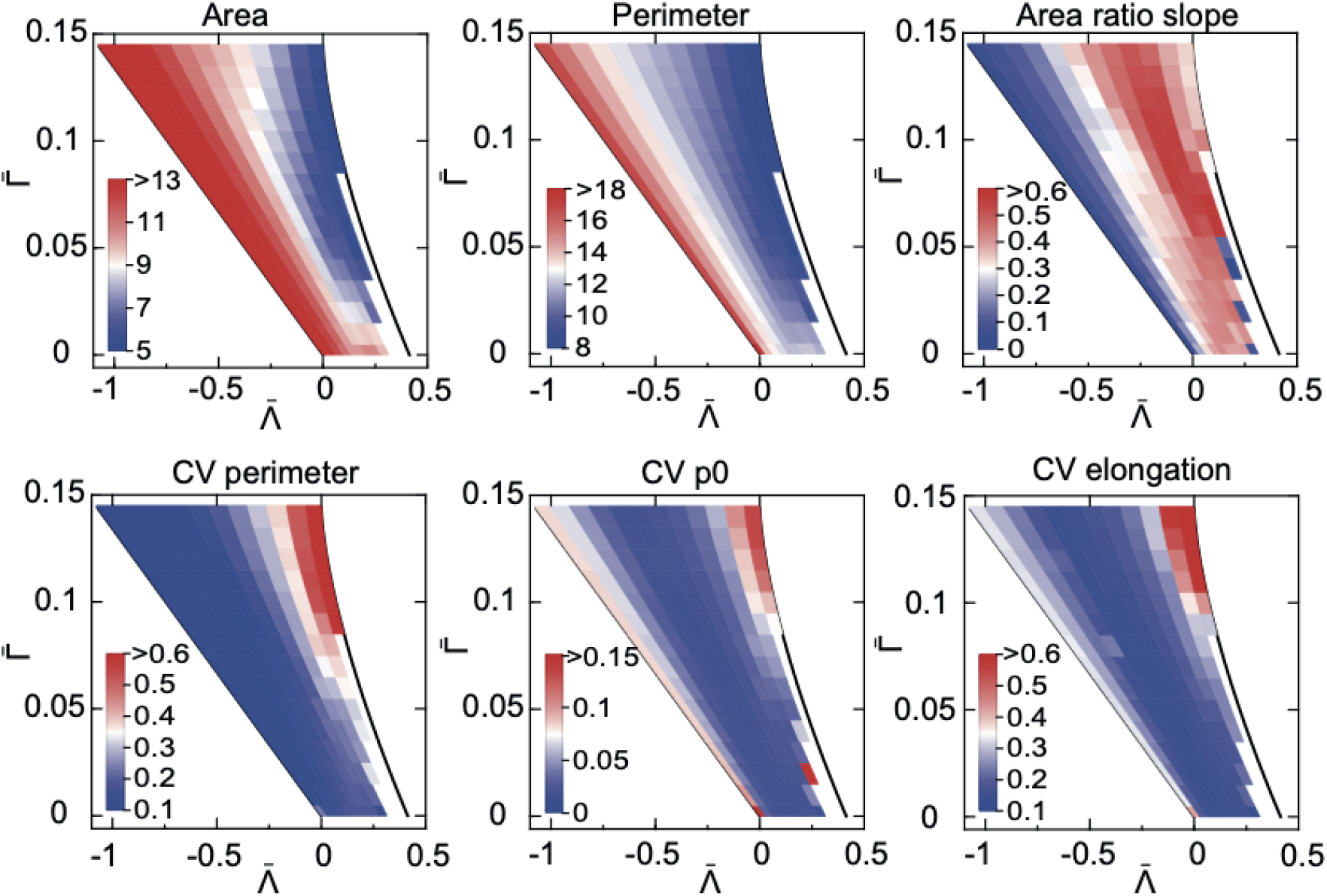
Cell shape descriptors in simulations. Mean value of cell shape descriptors (cell area, cell perimeter, area ratio slope, coefficient of variation of cell perimeter, coefficient of variation of shape index, coefficient of variation of cell elongation, as defined Table S3) for simulations performed at different values of 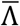 and 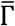 for proliferation rate *k*_*p*_ = 0.09h^-1^ and differentiation rate *k*_*n*_ = 0. The mean value was color-coded with scale indicated in insets of respective panels. The mean was estimated from all cells pooled together from 10 simulations for each pair of 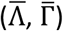 at the final time.

**Figure S6.**
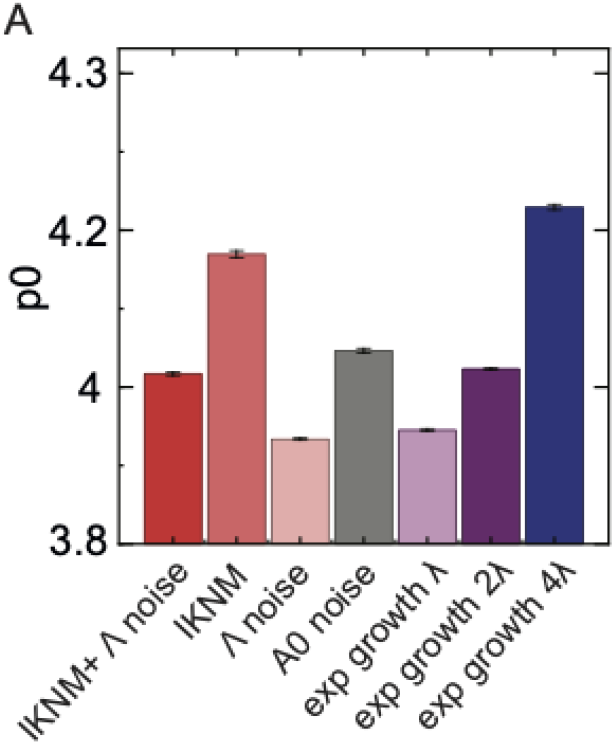
Cell shape index in simulations with different apical cell area kinetics during the cell cycle. The simulated conditions are as in Figure. 3E-F: IKNM+Λ noise (default condition), IKNM (no noise in line tension), Λ noise (cell divisions without IKNM, with noise in 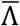), A0 noise (target area randomly drawn from a distribution), exp growth (exponential growth of the target area). See Methods for details. n=10 simulations per condition, region 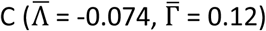.

**Figure S7.**
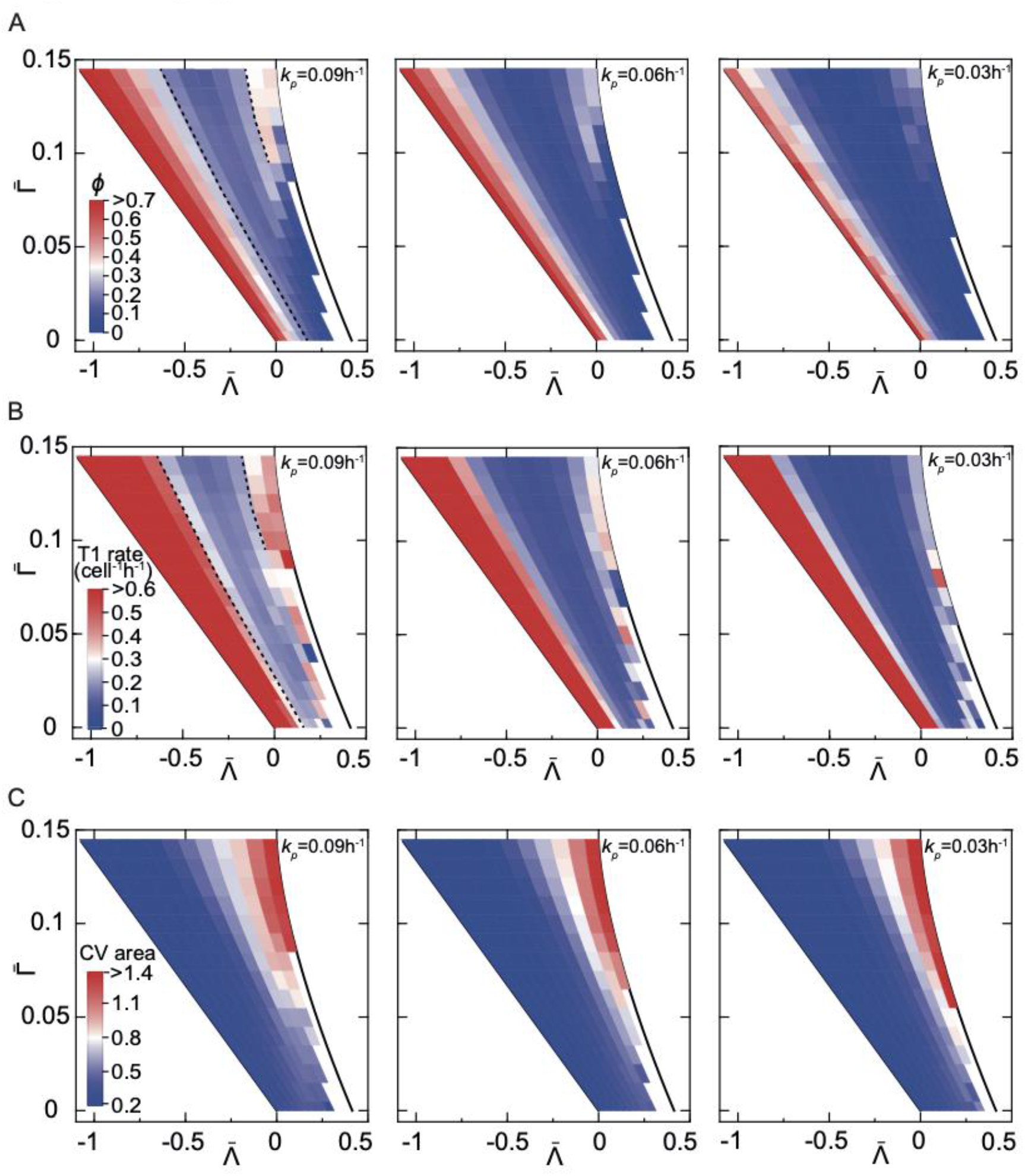
Fragmentation coefficient, T1 rate and cell area CV at different proliferation rates. **A**. Fragmentation coefficient at different values of 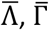for high (*k*_*p*_ = 0.09h^-1^), intermediate (*k*_*p*_ = 0.06h^-1^) and low (*k*_*p*_ = 0.03h^-1^) rates of proliferation. Differentiation rate *k*_*n*_ = 0. **B**. Rate of T1 transitions with the same set of parameters as in A. **C**. The coefficient of variation of cell areas with the same set of parameters as in A. The estimates are mean values from 10 simulations for each pair of 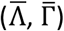 at the final time.

**Figure S8.**
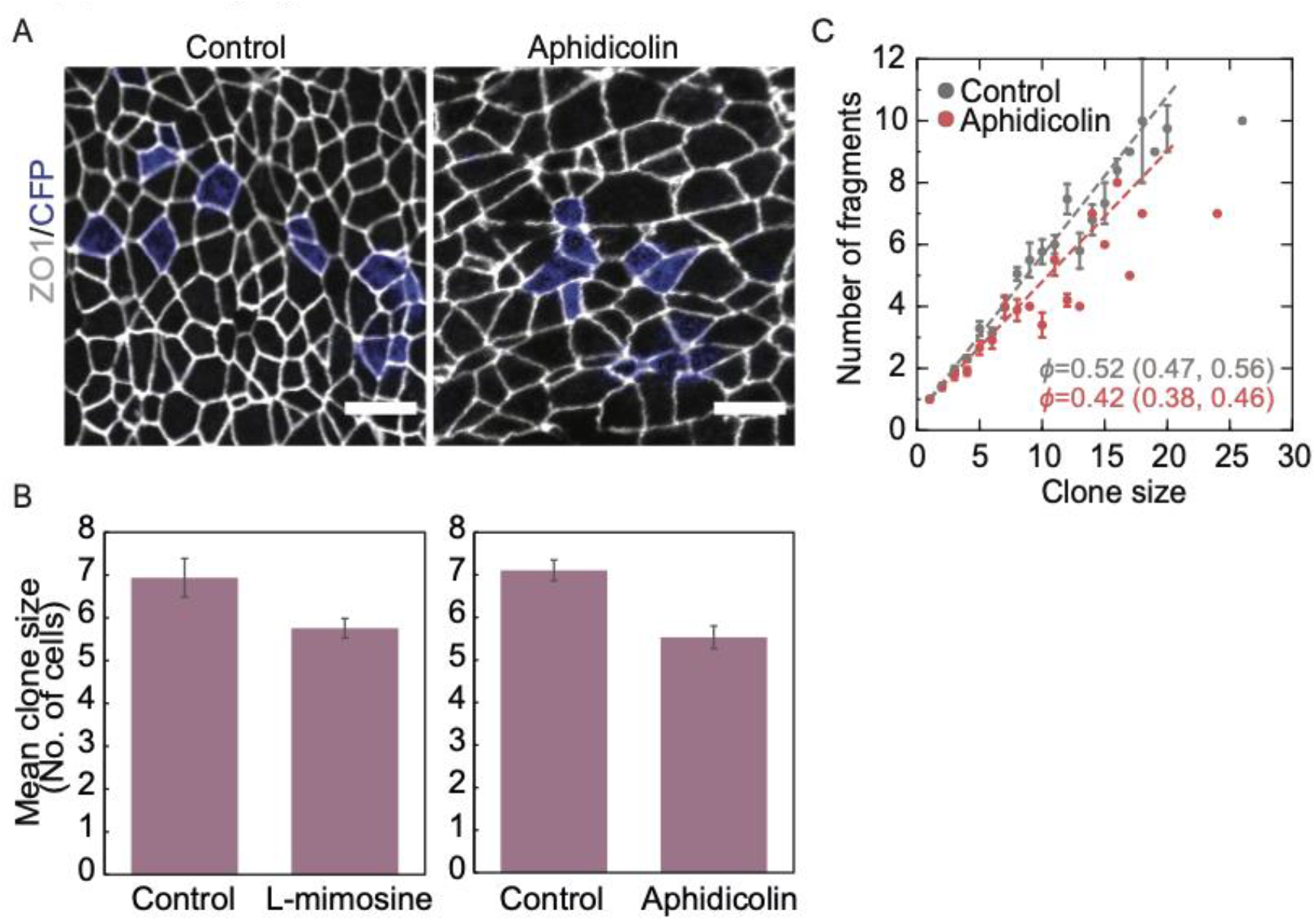
Inhibition of cell proliferation affects clone fragmentation and growth. **A**. Confetti clones (CFP) from embryos injected with tamoxifen at E7.5 and cultured ex utero from E8.5 for 42h with 800nM aphidicolin or control medium. Scale bars, 10 μm. **B**. Mean clone size for aphidicolin (A) and L-mimosine (Figure. 4B-C) conditions compared to control. **C**. Mean number of fragments per clone for a given clone size for control (n = 382 clones) vs aphidicolin treated embryos (n = 185 clones). Error bars, SEM.

**Figure S9.**
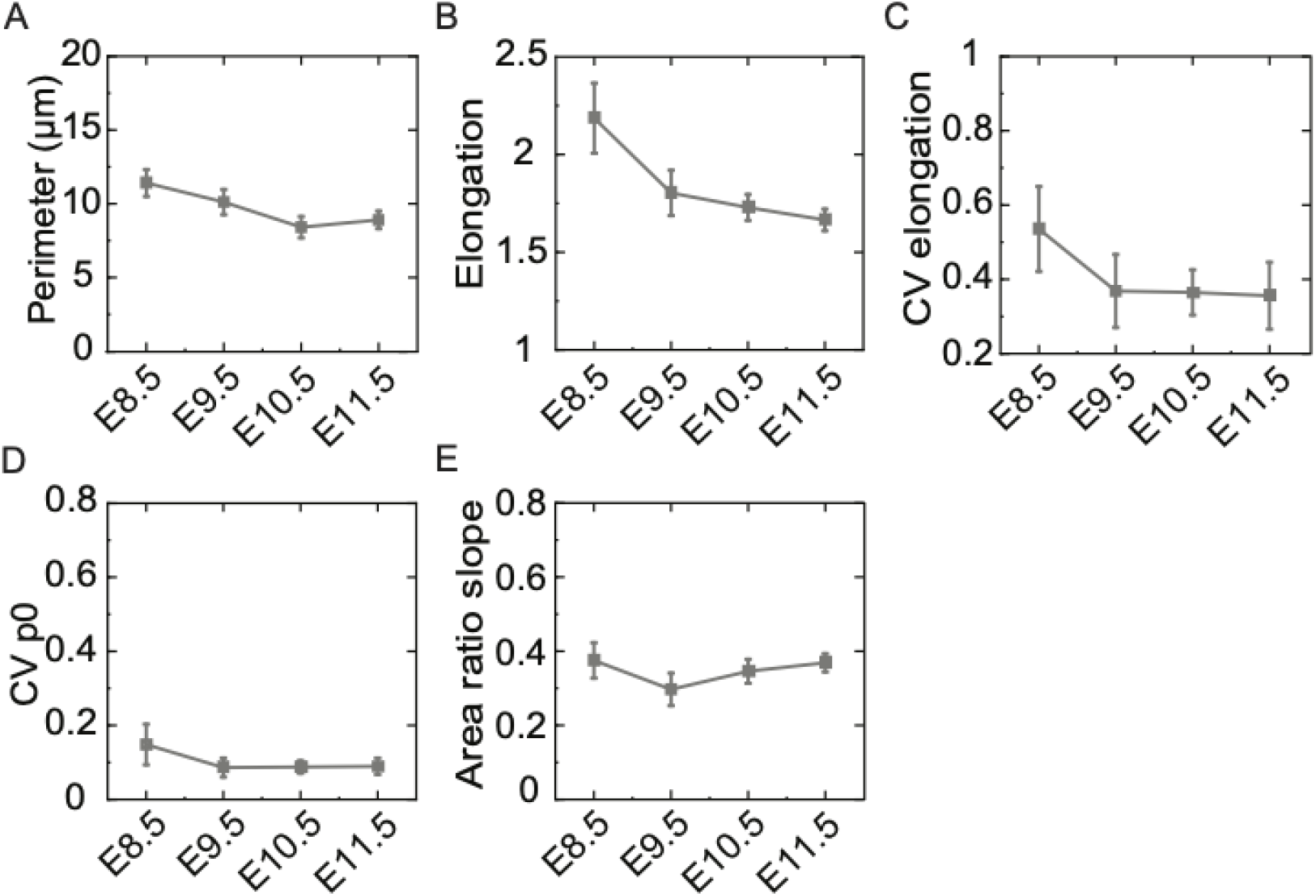
Cell shapes in neuroepithelia collected at different stages. **A**. Cell perimeter. **B**. Cell elongation. **C**. Coefficient of variation of cell elongation. **D**. Coefficient of variation of cell shape index. **E**. Area ratio slope. Mean values ± SEM are shown. Sample sizes as in Figure. 5B-E (see Table S1).

**Figure S10.**
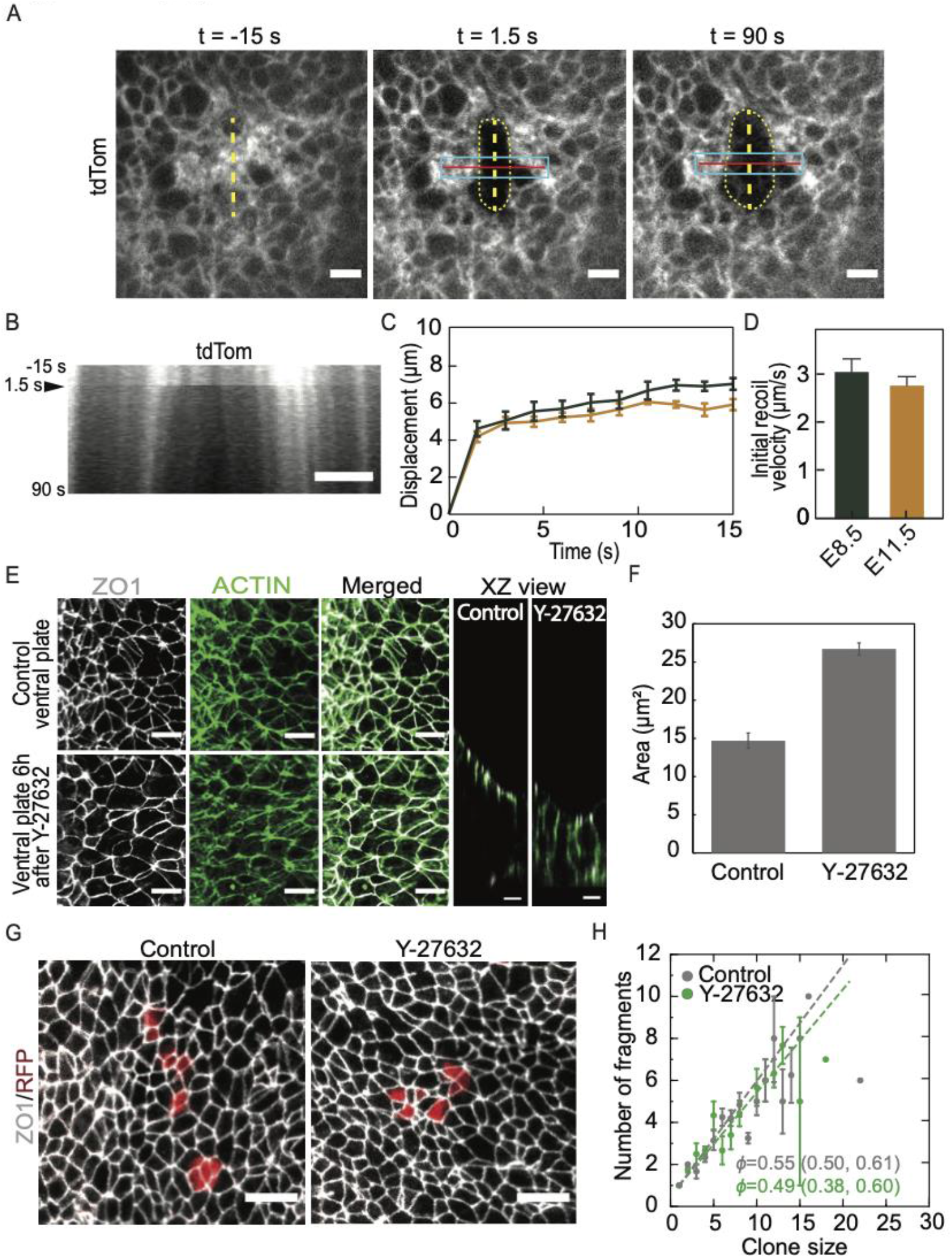
The neuroepithelium maintains its contractility and tension over time. **A**. Apical view of membrane tdTomato expressing neuroepithelium at E8.5 before and after laser ablation. Laser cut was performed at t = 0 s along the yellow dashed line. The resulting tissue displacement (yellow dotted line) relative to the cut line was quantified within the region outlined by cyan rectangle. **B**. Representative kymopgraph showing the change in fluorescence intensity along the red line in A. **C, D**. Mean tissue displacement over time (C) and initial recoil velocity ± SEM (D) after laser ablation at the indicated stages. Error bars, SEM. In D, Mann-Whitney test p-value>0.05. Sample size: n = 8 ablations from 4 embryos for E8.5, n= 5 ablations in 3 embryos at E11.5 **E**. Intermediate E8.5 neural plate stained for ZO1 (white) and actin (green) after treatment with 8μM Y-27632 or vehicle for 6 hours. First three panels, apical view, right-most panels – apicobasal view. The apical enrichment of actin is perturbed in inhibitor treated embryos. **F**. Mean apical cell area in E8.5 embryos treated with Y-27632 for 6h vs controls ± SEM. Double-sided T-test p<10^−9^. Number of cells analyzed: 1728 (Y-27632), 2922 (control). **G**. RFP labelled Confetti clones in Y-27632 treated embryos vs controls. Inhibitor treatment was performed for 6h at E8.5, subsequently embryos were cultured for 42h without inhibitor before harvesting for analysis. **H**. Fragmentation coefficient of control and Y-27632 clones (mean, 95% CI). Control n= 83 clones, Y-27632 n = 58 clones. Scale bars, 10 μm. Error bars, SEM.

## Notes

### Competing Interest Statement

The authors have declared no competing interest.

